# Ramularia leaf spot disease of barley is highly host genotype-dependent and suppressed by continuous drought stress in the field

**DOI:** 10.1101/2020.06.02.121491

**Authors:** Felix Hoheneder, Katharina Hofer, Jennifer Groth, Markus Herz, Michael Heß, Ralph Hückelhoven

## Abstract

Since the 1980s, Ramularia leaf spot (RLS) is an emerging barley disease world-wide. The control of RLS is increasingly aggravated by a recent decline in fungicide efficacy and a lack of RLS-resistant cultivars. Furthermore, climate change increases drought periods in Europe, enhances variable weather conditions and thus will have an impact on severity of plant diseases. Hence, identification of RLS-resistant cultivars and understanding of disease progression under abiotic stress are important aims in integrated disease management under climate change. In the present study, we evaluated quantitative RLS resistance of 15 spring barley genotypes under drought, controlled irrigation and field conditions between 2016 and 2019 and monitored microclimatic conditions within the canopy. We identified genotypes that show robust quantitative resistance to RLS in different field environments. Our findings suggest that long-lasting drought periods create unfavourable conditions for the disease and supports that the extent and duration of leaf wetness is a key factor for RLS epidemics.

## Introduction

Barley (*Hordeum vulgare* L.) is the fourth most important cereal crop plant worldwide, especially due to its robustness and adaptation to various and mostly restricted environments. On a global scale, about 85 % of barley production is utilised for animal feed, whereas a small proportion is used for human consumption (Fischbeck, 2002). Due to specific qualities of barley grain in favour of malting, it is commonly used for brewing beverages, which give high added values in the production and supply chain. Moreover, under wide span of climate zones and very different growing conditions, barley possesses high yield potential and can be harvested after a comparatively short vegetation period. Thus, importance of barley will likely grow under increasingly unfavourable environmental conditions (e.g. increase of heat and drought) caused by a changing climate (Newton et al., 2011). However, as recently reported, global malt production is sensitive to climate change, which severely may affect future prices of barley and malt products (Xie et al., 2018).

Additionally, climate change is further driving important diseases in cereal crops, e.g. cereal rust, Fusarium crown and foot rot (Luck et al., 2011). The shift of various climate factors like extreme and frequent precipitation events, drought or elevated mean temperature may directly and indirectly influence geographical appearance and distribution, seasonal phenology and further population dynamics of pathogens and pests (Chakraborty et al., 2000; Anderson et al., 2004; Juroszek & von Tiedemann, 2013). From a field-scale view, climate and local weather are determining factors, which variously affect microclimate within canopy, the dynamics of inoculum spreading and infection process of a certain pathogen and further disease severity (Chakraborty et al., 2000; Luck et al., 2011). Additionally, the simultaneous occurrence of multiple stress factors is particularly fatal for crop production. It triggers different molecular pathways in stress-signalling resulting in a synergistic or antagonistic crosstalk of stresses which affect growth and plant defence (Pandey et al., 2015; Ramegowda & Senthil-Kumar, 2015). Consequently, prediction about the occurrence and severity as well as management of emerging plant diseases (e.g. Ramularia Leaf Spot of barley) (Havis et al. 2015) is restricted and a challenge for crop production. Elevated risks are further aggravated by the predicted increase of variability in weather conditions and complex modifications in host-pathogen interactions, both driven by a changing climate system (Chakraborty et al., 2000; Atkinson & Urwin, 2012). Improvement of host-resistance and utilization of genetic resources are effective, sustainable and promising tools to meet such challenges in future cereal crop production by cultivation of environmental robust and healthy crops.

The fungus *Ramularia collo-cygni* (Sutton and Waller) is the causal biotic agent of the Ramularia Leaf Spot (RLS) disease of spring and winter barley (*Hordeum vulgare* L.). The mitosporic ascomycete was first isolated and described in Italy in 1893 as *Ophiocladium hordei* (Cavara, 1893) and later reclassified to *Ramularia collo-cygni* (Sutton & Waller, 1988). Since the 1980s, *R. collo-cygni* drew increasing attention in agriculture and research, when unspecific necrotic leaf symptoms were first associated as a fungal disease (Sachs, 2006). Since then, RLS is an emerging disease in barley causing increasingly economical relevant epidemics in several important barley growing countries in Europe and world-wide (Havis et al., 2015). Yield losses are attributed to a reduction of grain size. Average losses are recorded with 5-10 %, which can peak up to 75 % on occasion (Pinnschmidt & Jørgensen, 2009; McGrann & Havis, 2017). Besides barley crops, *Ramularia collo-cygni* also infects other cereals like wheat (*Triticum aestivum* L.), triticale (*Triticum* x *Secale*) (Sutton & Waller, 1988), oat (*Avena sativa* L.), rye (*Secale cereale* L.), maize (*Zea mays* L.) and a broad range of grasses and weeds, though with lower economic impact. The role of alternative hosts mediating as green bridge is still a matter of discussion, even though symptom formation is hardly conspicuous, except on barley (Huss, 2004; Frei et al., 2007; Huss, 2008; Huss & Miethbauer, 2010; Kaczmarek et al., 2017, Stam et al., 2019). Symptoms occur mainly on upper leaves of barley in the late season post flowering and cause rapid leaf senescence. Furthermore, the fungus also infects tissues like culms, lemma, palea and in particular awns (Huss, 2004). In contrast, the fungus asymptomatically colonizes the host plant during a relatively long endophytic stage in the early season. At a later point in time, the fungus induces lesions, leaf senescence and the formation of spores exclusively occurring on necrotic tissue (Walters et al., 2008; Kaczmarek et al., 2017). From this stage, *Ramularia collo-cygni* switches to a necrotrophy pathogen and becomes very dominant in the field. Important triggers for this switch in lifestyle are associated with the interaction of abiotic factors and host ontogenesis during generative phase of infected barley crops (Walters et al., 2008; Schützendübel et al., 2008; Newton et al., 2010). Thus, it can be assumed that an increase of plant stress situations driven by global warming will likely enhance RLS disease outbreaks.

RLS symptoms are reddish to brown necrotic spots surrounded by a chlorotic halo and confined by leaf veins. The spots mainly develop on the sunlight-exposed sides of the leaf blade in an unspecific, speckle-like pattern and resemble physiological leaf spots (PLS). The interaction of abiotic stress-factors, physiological imbalances in leaf tissue resulting in oxidative stress, cultivar dependent physiology and RLS cause together the leaf spot complex in barley. Both RLS and PLS components result in a reduction of green leaf area and reduce yield potential (Oxley et al., 2002; Sachs, 2006; Heß et al., 2007; Schützendübel et al., 2008; Brown et al., 2014). Due to the fact that RLS can easily be confused with PLS and early small-sized symptoms caused by other necrotizing pathogens i.e. *Pyrenophora teres* (net blotch), particularly PCR-based diagnostic methods have been developed and contribute to a clear RLS/PLS differentiation, *R. collo-cygni* detection and fungal DNA quantification (Havis et al., 2006; Frei et al., 2007; Taylor et al., 2010). Both, RLS symptom formation and PLS directly coincide with maturation and further plant stress late in the season. In this context, the antioxidative defence system protects leaf tissue from an overload in reactive oxygen species (ROS) (Wu & von Tiedemann, 2004; Schützendübel et al., 2008). Under normal conditions, ROS are involved in metabolism, plant defence signalling, aging and whole plant ontogenesis (Apel & Hirt, 2004), but during plant ripening, an imbalance in the antioxidative system is associated with the fungal production of light-dependent and photodynamic rubellins A, B, C and D, released into the apoplast of infected barley leaves. Together with the secretion of other toxic compounds and cell wall degrading enzymes (Sjokvist et al., 2019), rubbellins may cause a relatively delayed symptom formation (Heiser et al., 2003; Miethbauer et al., 2003; Miethbauer et al., 2006).

Practices of Integrated Pest Management (IPM) including crop rotation, cultivation techniques (ploughing, support of decomposition of crop debris), disease monitoring and the choice of resistant varieties do not ensure a sufficient RLS control (Havis et al., 2018b). Diverse studies suggested that full RLS resistance is absent from the barley gene pool and rather moderate susceptibility is what can be observed in comparative genotype surveys (Reitan & Salamati, 2007; Havis et al. 2015). Several studies revealed inconsistent results on a possible linkage of the *mlo* resistance locus effective against powdery mildew in barley (*Blumeria graminis* f.sp. *hordei*) and the level of susceptibility to *Ramularia collo-cygni* and abiotic leaf spots (PLS) (Makepeace et al., 2007; Pinnschmidt & Sindberg, 2007; McGrann et al., 2014; Hofer et al., 2015). In this context, it is also questioned whether the extended cultivation of mildew resistant *mlo* barley lines since the mid to late 1980s coincide with the markedly emergence of RLS and the PLS in all important barley growing regions in Europe (Makepeace et al., 2007; Havis et al., 2015). However, *mlo* mediated resistance against powdery mildew of barley is not employed in winter barley, whereas winter barley is similarly susceptible to *R. collo-cygni* (Jørgensen, 1992; Makepeace et al., 2007; Czembor et al., 2016). Hence, *mlo* alone does not explain susceptibility of barley crops when compared to other small grain cereals.

Because fully resistant cultivars are not available, the control of RLS in barley is primarily based on chemical control by fungicides. A decline in fungicide efficacy up to a broad field resistance was recently observed in several European barley growing countries, concerning to major groups of fungicides: Thus, the control of RLS by quinone outside inhibitors (QoI), including strobilurines (Fountaine & Fraaije, 2009; Matusinksy et al., 2010), demethylation inhibitors (DMI) and succinate dehydrogenase inhibitors (SDHI) (Piotrowska et al., 2017; Rehfuß et al., 2019) is hardly effective anymore. The multi-site inhibitor chlorothalonil is solely a very effective chemical agent against RLS in cultivation of barley (Havis et al., 2018a). Nevertheless, the European Food Safety Association (EFSA) banned the use of chlorothalonil based fungicide products from 2020 onwards due to environmental safety concerns. Consequently, from the view of practical farming, effective management of RLS has to rely on alternative measures and disease resistance moves even more into the focus of an integrated approach.

The present study indicates the influence of late-terminal drought stress on the occurrence and severity of RLS in an assortment of 15 spring barley cultivars and current breeding lines, respectively. We used a rainout shelter to expose spring barley to drought stress under field conditions. Visual assessments of RLS, foliar diseases and the sporulation in the field reveal a general suppression of RLS under long lasting dry conditions. Furthermore, the determination of fungal DNA confirmed observations in the field. The data depicts great differences in RLS susceptibility between spring barley genotypes and identifies candidates for breeding RLS-resistant cultivars. We discuss potential effects of climate change along with drought conditions on RLS disease in Central Europe. The results support the understanding of the complex interaction between plant genotypes under stress conditions and the pathogenesis of RLS.

## Material and methods

### Field trials under controlled drought stress

For studies about the occurrence of RLS disease on drought stressed spring barley under field conditions, a rainout shelter was used (moveable transparent polyvinyl roof with open fronts, size: approximately 12 x 34 m) at the Bavarian State Institute of Agriculture (LfL) in Freising (Southern Germany; soil: loamy-silt). The rainout shelter was equipped with a rain and wind sensor, to shield the field from rainfalls. 59 genotypes were sown twice in four blocks of randomized microplots (two rows with a length of 1 m) in 2016 and 2017.

The blocks were regularly irrigated with 20 mm of water once a week by a permanently installed irrigation system 2 m above ground. The amount of irrigation was set according to long-time rainfall at the location in Freising. During GS 50 (approximately at the beginning of May), irrigation of two cater-cornered blocks was stopped to gain permanent drought stress (soil moisture tension below −500 hPa) from the beginning of spike emergence to full maturing. The control plots were continuously irrigated once a week. During periods of heat, the control plots were irrigated twice a week to maintain sufficient water supply as required. Soil moisture was monitored with tensiometers in depths of 15 cm, 45 cm and 75 cm three times per week between begin of May and mid of July. Exemplary data of soil moisture tension in a depth of 45 cm is represented in figure S-1.

Agronomic data of the 59 genotypes is provided in table S-1. The collection showed great variation concerning all determined traits. Significant differences were also found between the non-irrigated plots and the control plots for most traits.

In season 2018, the assortment was narrowed down to 15 candidate genotypes showing variability in RLS resistance and furthermore stable performance and valuable agronomic traits under drought conditions in the two previous seasons (table S-1, highlighted genotypes). Due to a better comparability, only two-rowed and adapted genotypes combined with other favourable traits were selected. For example, genotypes “13/594/74”, “Barke”, “IPZ 24727” and “Marnie” had no reduced kernel size under drought stress. “RGT Planet” and “STRG 706/16” were characterized by a high grain weight per double row in the non-irrigated plots. Genotypes with a high number of ear carrying stalks were “Argentinische DH 168”, B0004”, “Barke”, “RGT Planet” and “STRG 706/16”. Plants were grown in four randomized blocks consisting one field plot of each cultivar. Each plot had a size of 3.5 square meters (1.5 x 2.3 m). The treatments “non-irrigated” and “control” were performed as described above.

For reasons of consistency, only results of the 15 candidate genotypes for all the seasons are presented throughout the paper.

The plants were cover-sprayed with 1.5 l/ha of the fungicide Input^®^ Xpro (Bayer Crop Science) approximately at growth stage 39 to control commonly occurring foliar diseases. Fertilization, insecticides and herbicides were used according to standard agricultural practises. After full ripening (GS 99), micro plots were harvested and threshed manually, macro plots were harvested with a single plot combine harvester. Sowing dates, recorded growth stages and dates of harvest of each year are listed in table S-2 (supplements).

### Additional field trials

A second location at Weihenstephan (Freising, South Germany; soil: loamy-silt), about 1.5 km away from the rainout shelter was used to assess natural infection of *R. collo-cygni* and RLS disease progression. Therefore, 59 spring barley genotypes were sown in microplots (two rows of plants with a length of 1 m) in 2016 and 2017. In the seasons 2018 and 2019, 15 candidate genotypes (table S-1) of the assortment were sown in standard field plots (size 1.5 x 7.5 m), respectively.

Plants were cover-sprayed once with 0.8 l/ha of the fungicide Prosaro^®^ (Bayer Crop Science) at GS 32 to control other foliar diseases except *Ramularia collo-cygni*. Nitrogen fertilizer and herbicides were used according to standard agricultural practise. After full ripening (GS 99), micro plots were harvested and threshed manually, macro plots were harvested with a plot combine harvester. Dates of sowing, recorded growth stages and harvest of each year are listed in table S-2 (supplements).

For the estimation of yield and stability of performance, the entire assortment was planted in macro plots ranging from 6.3 to 11.3 square meters in six additional environments (three locations in Bavaria, one in Lower Saxony, one in Saxony-Anhalt and one in France) in the season 2018. Plants were grown in two randomized blocks at each location. Fertilization and pesticide treatments were performed according to local agricultural practises. As mentioned above, only the results of the 15 genotypes of the narrowed assortment are presented and discussed in this paper.

### Weather conditions and microclimate within plant canopy

Average temperature, sum of precipitation and leaf wetness were recorded by weather station in Freising during the seasons 2016 to 2019. The data was assessed from the agro-meteorology web portal of the Bavarian State Institute of Agriculture (Agrarmeteorologie Bayern, 2020). In addition, data logger (HOBO^®^ Pro v2 data logger, Onset, MA, USA) were placed within the plant canopy to measure on-site temperature and relative humidity and to automatically calculate the temperature of the dew point in a time interval of one hour. Data logger were read out with HOBOware Pro software system (Onset, MA, USA). Mean values were calculated for each week during the growing season.

### Assessment of symptoms or sporulation and leaf sampling

To examine foliar disease infestation with focus on Ramularia leaf spot and physiological leaf spots, a visual assessment of all common foliar diseases of barley was conducted at early to mid-dough stage (growth stage 83-85). Physiological leaf spots (PLS) and Ramularia leaf spots (RLS) were summed up as a leaf spot complex due to unspecific differentiation in the field. For accurate determination of RLS and sporulation, up to 10 flag leaves (F) and the leaf stage below (F-1) were randomly sampled from each cultivar at growth stage 83 to 85 in both trials. Leaves were stored at −20 °C until further processing.

Sporulation was assessed by the use of a binocular microscope at a maximum magnification of 40 and bright light from side in a flat angle. Therefore, clusters of sporangiophores on the bottom side of several leaves were examined. Sporulation was determined as mean leaf area with visible sporangiophores in percent.

### Isolation of genomic DNA from leaf material

Genomic DNA was extracted according to the protocol of Fraaije et al. (1999) including minor modifications as published by Hofer et al. (2016). DNA concentration (ng total DNA μl^-1^) and quality were measured by the use of a NanoDrop^®^ ND-1000 spectrophotometer (Thermo Fisher Scientific, MA, USA). In a final step, the samples were adjusted to a concentration of 20 ng total DNA μl^-1^ with sterile double-distilled water. Extracted DNA samples were stored at 4 °C until further processing.

### Quantification of *R. collo-cygni* DNA

*R. collo-cygni* DNA was quantified by a real-time quantitative polymerase chain reaction (qPCR) following a Taq-Man qPCR protocol according to Taylor et al. (2010). The PCR amplification was performed by the use of an MX3000P Multiplex Quantitative PCR System (Stratagene, CA, USA). Data analysis was carried out with MxPRO qPCR software (Stratagene, CA, USA).

### Calculation of disease severity ranking

To evaluate the resistance to RLS in manner, which balances year-effects of disease severity, a ranking list was calculated. Therefore, the mean ranking of each assessed disease parameter (mean ranking for symptomatology, sporulation and DNA contents) and a total mean rank was calculated for each genotype and year (minimum rank: 1; maximum rank 15) and then averaged over years. Low ranks indicate resistance; high ranks indicate susceptibility. Shared ranks were scored with same single value.

### Stability analysis

In order to determine the stability of genotypes according to yield parameters across environments the additive main effect and multiplicative interaction (AMMI) model was used. It fits the additive effects for genotypes and environments and multiplicative terms for genotype x environment interactions (Crossa et al. 1990). AMMI stability values (ASV) were calculated for each of the 15 genotypes according to Purchase et al. (2000) to rank genotypes according to trait stability. The lower the ASV, the more stable is a genotype. The calculated yield parameter stability index (YPSI) is based on summing the ranking of overall mean performances for each trait and the ranking for ASV for each trait. The YPSI was calculated as follows:

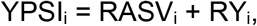

where YPSI_i_: yield parameter stability index for the i^th^ genotype across seasons/treatments; RASV_i_: rank of the i^th^ genotype across seasons/treatments based on ASV; RY_i_: rank of the i^th^ genotype based on mean performance across seasons/treatments. Beside of yield, the traits of interest were fraction of kernels with a size > 2.5 mm, fraction of kernels with a size > 2.8 mm and thousand kernel weight (TKW). The yield parameters were assessed for the 15 genotypes in the rainout shelter experiment, our own field trial and six additional field locations in 2018. YPSI ranks of each genotype were summed for all traits and the genotype with the smallest rank sum was considered to be the best across traits.

The AMMI analysis as well as the stability values (ASV) and indices (YPSI) were calculated using R version 3.5.2. (www.r-project.org) and the package *agricolae* (Mendiburu & Simon, 2015).

## Results

### Evaluation of Ramularia resistance under irrigated and drought conditions

For determination of basal resistance to RLS of several spring barley genotypes, we assessed disease severity after natural infection by three diagnostic tools on samples from either the field or from a rainout shelter. (i) We rated the leaf area with RLS symptoms and plant stress related PLS together (RLS+PLS), because symptoms are difficult to differentiate without microscopy. Therefore, the data illustrates the Ramularia leaf spot complex of barley. (ii) In parallel to visual assessment, leaf samples (flag and flag-1 leaf) were collected to examine leaf area with sporulation of *Ramularia collo-cygni* with the help of a stereo microscope. (iii) Additionally, genomic DNA was extracted from flag and flag-1 leaf samples to assess the development of *R. collo-cygni* in upper leaves [Fig. 1a, 1c, 1e]. We used an assortment of 15 two-rowed spring barley genotypes, which was preselected for high and stable yield, homogenous plant height, similar heading date and high performance in different field environments and under drought stress (table S-1).

**Figure 1.**
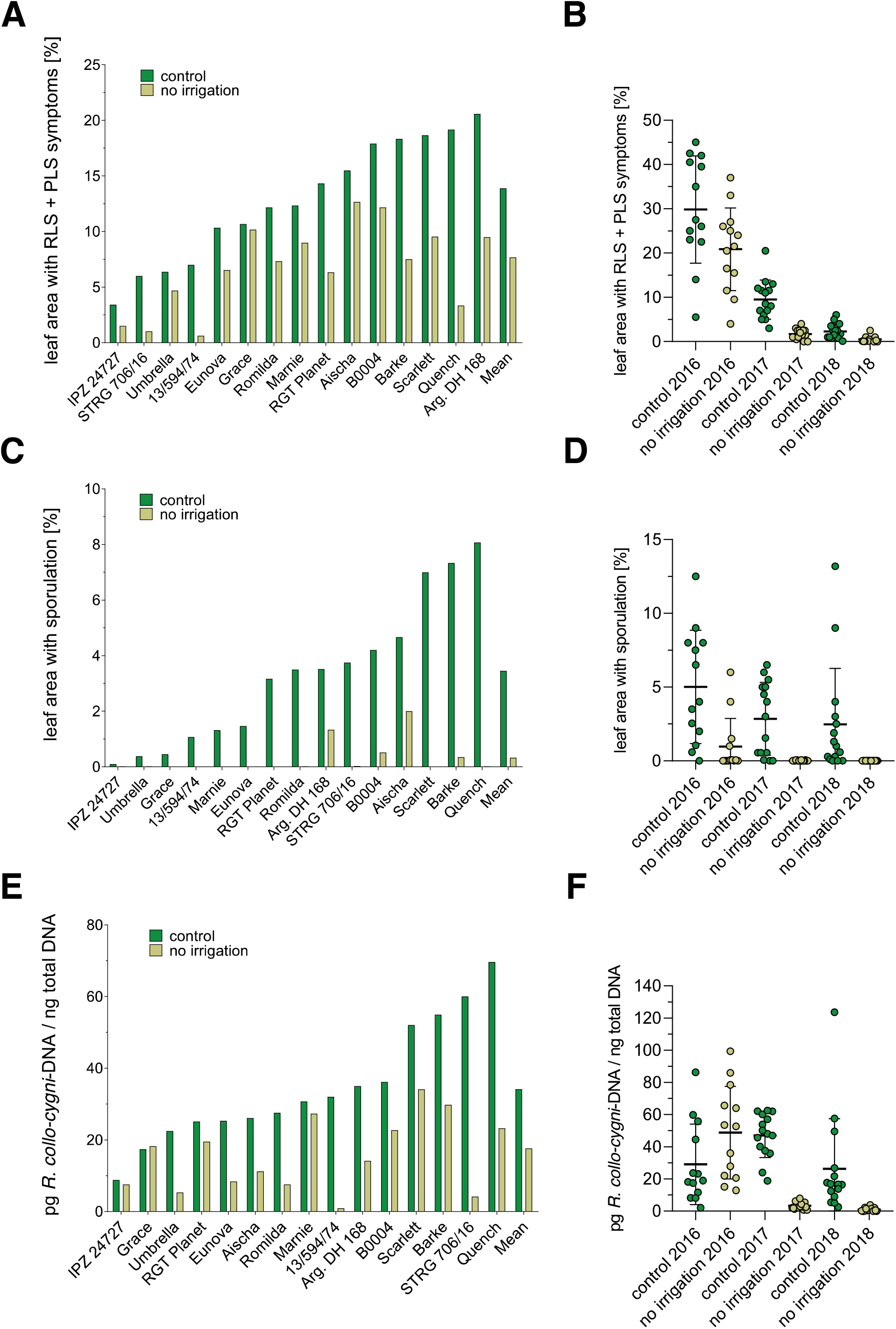
Levels of Ramularia leaf spot infestation according to assessed disease parameters under drought stress and irrigated control in the rainout shelter experiment between 2016 and 2018. The bars indicate calculated mean values of three years. [A] represents percentage of leaf area with Ramularia and physiological leaf spots. [C] indicates percentage of leaf area with sporulation. [E] shows detected values of Ramularia-DNA in pg Rcc DNA / ng total DNA. [B], [D] and [F] show single values and the total mean of each disease parameter in the control and under drought stress for each season. Error bars indicate the standard deviation of the mean.

In the rainout shelter, we compared irrigated plots from those under permanent drought stress from GS 50 onwards over three consecutive years from 2016-2018. The genotypes growing under irrigation showed a strong differentiation in RLS severity. Noticeable symptoms were found on upper leaves, which also reflected fungal DNA contents and sporulation. Under drought, all tested 15 genotypes were less severely infected with *R. collo-cygni* and showed less RLS symptoms when compared to the irrigated control plots. RLS was in general the dominating foliar disease in the control and under drought [Fig. S-2]. The results revealed a general decrease of Ramularia disease after application of drought stress between spike emergence and full ripening stage over three consecutive years: symptomatology, sporulation and DNA-contents in the upper leaves were largely corresponding to each other [Fig. 1a – 1f, Figure S-3]. A Pearson’s correlation revealed significant relations between leaf area with sporulation and with Ramularia-DNA and leaf area with sporulation and RLS symptoms under irrigation. RLS symptoms significantly correlated with Ramularia-DNA contents and leaf area with sporulation under drought conditions [Figure S-3].

Mean levels of symptomatic leaf area ranged between 3.4 % (breeding line IPZ 24727) and 20.5 % (Argentinische DH 168) in the irrigated control. Under drought stress, the highest mean of symptomatic leaf area was 12.6 % (Aischa), the lowest recorded mean area was 0.6 % (breeding line 13/594/74). Symptomatic leaf area of cultivar Grace was almost equal in the irrigated control compared to plants growing under drought stress (control: 10.7 %; drought stress: 10.2 %) [Fig. 1a].

Leaf area with visible *R. collo-cygni* sporulation did not exceed 8.1 % in the irrigated controls. The genotypes Quench, Barke and Scarlett showed the highest rates of sporulation (8.1 %, 7.3 % and 7.0 %). Little sporulation (< 1 %) was found for the genotypes IPZ 24727, Umbrella and Grace, followed by breeding line 13/594/74 (1.1 %), Marnie (1.3 %) and Eunova (1.5 %) [Fig. 1c].

Under drought stress conditions, we detected sporulation only on 9 out of 15 genotypes. Mean values remained below 2.0 % (Aischa). On leaves of Grace, RGT Planet, Marnie, Scarlett, Eunova and 13/594/74, no sporulation was recorded under drought conditions [Fig. 1c].

Contents of Ramularia-DNA differed among the tested genotypes in a range between 8.5 and 87.3 pg/ng total DNA (factor of 10.3) in the irrigated control. Under these conditions, highest average DNA content (69.6 pg/ng total DNA) was found in leaves of Quench and the lowest average content (8.8 pg/ng total DNA) in IPZ 24727 [Fig. 1e].

Mean DNA contents of drought-stressed genotypes remained below the contents in the corresponding irrigated control, except for Grace (control 17.4 pg/ng total DNA; drought stress: 18.3 pg/ng total DNA). DNA contents ranged between 0.9 pg/ng total DNA for 13/594/74 and 34.1 pg/ng total DNA in Scarlett under drought conditions. The most noticeable differences of fungal DNA contents between control and drought stress accounted for the breeding lines STRG 706/16 (control: 60.0 pg/ng total DNA; drought stress: 4.2 pg/ng total DNA) and 13/594/74 (control: 32.0 pg/ng total DNA; drought stress 0.9 pg/ng total DNA) [Fig. 1e].

A comparison of single years revealed strong variations in RLS severity in the rainout shelter over the three years [Fig. 1b, 1d, 1f]. RLS and PLS symptoms predominantly occurred in 2016 (total mean, control: 29.8 %; total mean, drought conditions: 20.8 %). In 2017, total mean leaf symptoms did not exceed 9.5 % (control) and 1.7 % (drought conditions), respectively and remained below 2.3 % in the control in 2018 [Fig. 1b]. Variation in total mean of leaf area with sporulation was comparable to leaf symptoms across single years. Total mean leaf area with sporulation was highest in the control in 2016 (5.0 %), followed by 2017 (2.9 %) and 2018 (2.5 %). Under drought conditions, sporulation was only detected in 2016, where mean sporulating leaf area reached 1 % [Fig. 1d]. Mean Ramularia-DNA contents in upper leafs of plants under irrigation were highest in 2017 (47 pg/ng total DNA), followed by 29.1 pg/ng total DNA in 2016 and 26.3 pg/ng total DNA in 2018. The highest mean DNA content was measured in upper leafs under drought conditions in 2016 (48.8 pg/ng total DNA), whereas DNA contents in 2017 (mean: 3.3 pg/ng total DNA) and 2018 (mean: 1.0 pg/ng total DNA) remained at a low level [Fig. 1f].

To rate resistance to RLS and to balance high variations in disease severity over the three years, we calculated ranks for each disease parameter and genotype (rank 1 to 15 for individual genotypes and years). From that, the total average rank over three years and three disease parameters (n = 9) was calculated and is presented in figure 2. For irrigated conditions, the IPZ 24727 (mean rank 2.2), Umbrella (mean rank 3.0) and Grace (mean rank 4.0) ranked lowest and hence they can be rated as quantitatively resistant to RLS under the given conditions. IPZ 24727 (mean rank 3.2) and Umbrella (rank 4.1) also showed high resistance under drought stress. Quench (mean rank 10.9), Barke (mean rank 10.8), breeding line STRG 706/16 (mean rank 10.6) and Scarlett (mean rank 10.4) were the most susceptible cultivars in the irrigated controls. IPZ 24727 and breeding line 13/594/74 showed lowest disease parameters under drought stress (average ranking over three years: 3.2 and 3.3), but breeding line 13/594/74 was less resistant in the irrigated control (mean rank 7.0). The breeding line B0004 and the genotype Argentinische DH 168 ranked highest in terms of RLS severity under drought stress.

**Figure 2.**
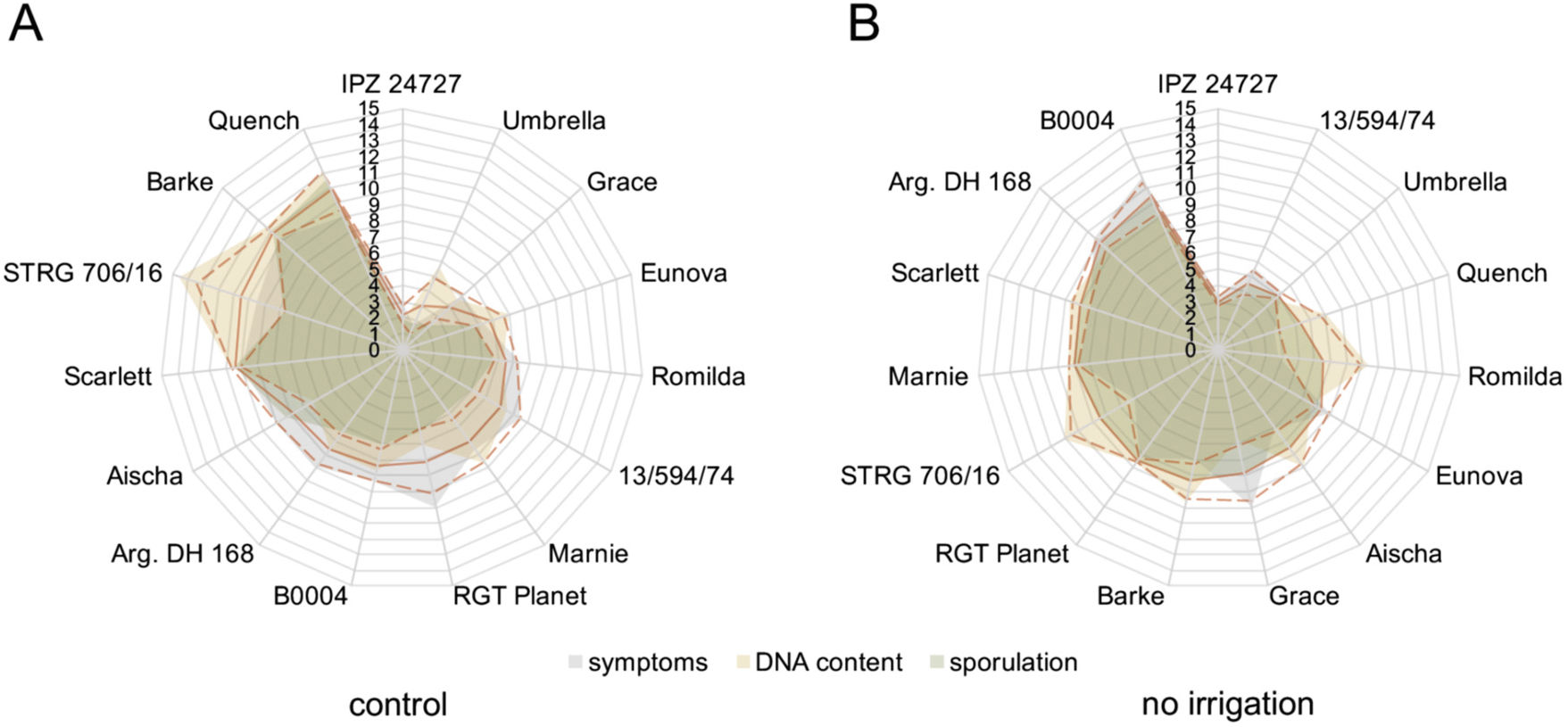
Disease severity ranking according to assessed disease parameters in the irrigated control [A] and under drought stress [B] in the rainout shelter experiment between 2016 and 2018. The radar plots show mean ranks over the three years for each disease parameter. The red line indicates the total mean rank over three years and three disease parameters (n = 9 ranks per genotype); dashed red lines indicate positive and negative standard deviation of the mean. The genotypes are sorted clockwise by total mean rank whereby low ranks indicate quantitative resistance and high ranks indicate higher susceptibility to RLS. Individual ranks are in a range of minimum 1 and maximum 15. If two or more lines share the same rank, the same value was associated to those lines.

Taken together, the data reveal a general decrease of *Ramularia collo-cygni* infection rates and suppression of sporulation in the drought-stressed barley plants compared to the control. Moreover, the assortment of cultivars and current breeding material showed strong differentiation in the level of quantitative resistance against RLS. Differentiation of RLS resistance was more pronounced under irrigation when compared to non-irrigated conditions. Particularly, the genotypes IPZ 24727 and Umbrella were little affected by RLS disease under both conditions.

### Monitoring of microclimatic conditions under drought and irrigation

We used data loggers to measure average temperature, average relative air humidity and the dew point temperature within the canopy for investigation of effects of drought on microclimate. Measurements were conducted between first week of June and fourth week of July in 2017 and 2018. The data is represented as weekly average [Tab. 1].

**Table 1.** Recorded microclimate of drought stressed (no irrigation) spring barley and in the weekly irrigated control plots in the rainout shelter experiment in 2017 and 2018. Data was collected by one data logger for each treatment (no irrigation, control) placed in a radiation shield within the canopy on the ground. The table shows weekly mean temperature and mean relative humidity between the 1^st^ week of June and 4^th^ week of July. The temperature of dew point was automatically determined by the data logger based on temperature and relative humidity. Measurements were conducted in time frames according to late growth stages (booting to grain maturity).

|  | temperature [°C] |  |  |  | relative humidity [%] |  |  |  | dew point [°C] |  |  |  |
| --- | --- | --- | --- | --- | --- | --- | --- | --- | --- | --- | --- | --- |
|  | 2017 |  | 2018 |  | 2017 |  | 2018 |  | 2017 |  | 2018 |  |
|  | control | no irrigation | control | no irrigation | control | no irrigation | control | no irrigation | control | no irrigation | control | no irrigation |
| 1<sup>st</sup> week of June | 20.9 | 21.9 | 20.7 | 23.2 | 78.3 | 66.4 | 84.0 | 67.1 | 16.5 | 14.7 | 17.6 | 16.1 |
| 2<sup>nd</sup> week of June | 15.9 | 16.9 | 21.0 | 23.9 | 89.0 | 73.8 | 87.9 | 68.7 | 13.9 | 11.8 | 18.6 | 17.1 |
| 3<sup>rd</sup> week of June | 19.0 | 19.9 | 17.7 | 19.9 | 82.1 | 68.3 | 86.6 | 68.9 | 15.6 | 13.3 | 15.1 | 13.3 |
| 4<sup>th</sup> week of June | 21.2 | 23.1 | 17.3 | 19.8 | 82.4 | 65.9 | 83.5 | 62.0 | 17.7 | 15.6 | 14.2 | 11.8 |
| 1<sup>st</sup> week of July | 19.9 | 21.7 | 17.8 | 20.2 | 85.3 | 69.4 | 80.2 | 63.3 | 17.1 | 15.3 | 13.8 | 11.9 |
| 2<sup>nd</sup> week of July | 20.7 | 23.0 | 19.8 | 21.9 | 81.9 | 63.0 | 80.0 | 63.0 | 17.2 | 14.9 | 15.7 | 13.5 |
| 3<sup>rd</sup> week of July | 20.1 | 21.0 | 19.1 | 20.6 | 79.0 | 68.4 | 78.0 | 65.1 | 15.9 | 14.5 | 14.7 | 13.1 |
| 4<sup>th</sup> week of July | 23.0 | 23.8 | 21.2 | 22.6 | 65.7 | 57.9 | 75.8 | 66.3 | 15.4 | 14.0 | 16.0 | 14.9 |
| **mean** | **20.1** | **21.4** | **19.3** | **21.5** | **80.5** | **66.6** | **82.0** | **65.5** | **16.2** | **14.2** | **15.7** | **13.9** |

During growth period in 2017, the mean temperature under dry conditions was increased by 1.3 °C compared to the irrigated control. At several points in time, the difference in mean temperature exceeded a range of 0.8 to 1.9 °C in 2017 between anthesis (mid of June) to full ripening (third to fourth week of July). Between the first week of June (GS 50: spike emergence) and third week of July (GS 99: grain maturity) in 2018, the drought-induced differences in mean temperature was between 1.4 and 2.9 °C. In both seasons, increased average temperature was accompanied by a lower relative air humidity in the canopy without irrigation. Total average air humidity in the irrigated plots was about 80 % (80.5 %, 82.0 %) in both seasons during the recorded period. The difference in average relative air humidity amounted to 13.8 % (2017) and 16.4 % (2018) between irrigated control and drought conditions respectively. Accordingly, the total mean dew point temperature was higher in the irrigated control by 1.9 °C in 2017 and by 1.8 °C in 2018, respectively.

### Monitoring of leaf wetness in the rainout shelter experiment

In addition to microclimate measurements, leaf wetness was monitored every 10 minutes in the canopy of the irrigated control and in the non-watered plots in the season of 2017 [Fig. 3]. The data was further used to calculate daily average leaf wetness. During the monitored time frame, the mean leaf wetness was significantly higher in the control (60.0 %) than in the non-irrigated plots (14.2 %) [Fig. 3a]. The progression of leaf wetness revealed higher fluctuating average leaf wetness in the irrigated plots (18.8 % - 100.0 %) compared to the plants growing under drought (3.8 % - 38.4 %). Furthermore, leaf wetness in the non-watered plots never reached the level that was simultaneously recorded in control plots [Fig. 3b]. This shows that the applied drought conditions strongly influenced the microclimate in the canopy and leaf wetness.

**Figure 3.**
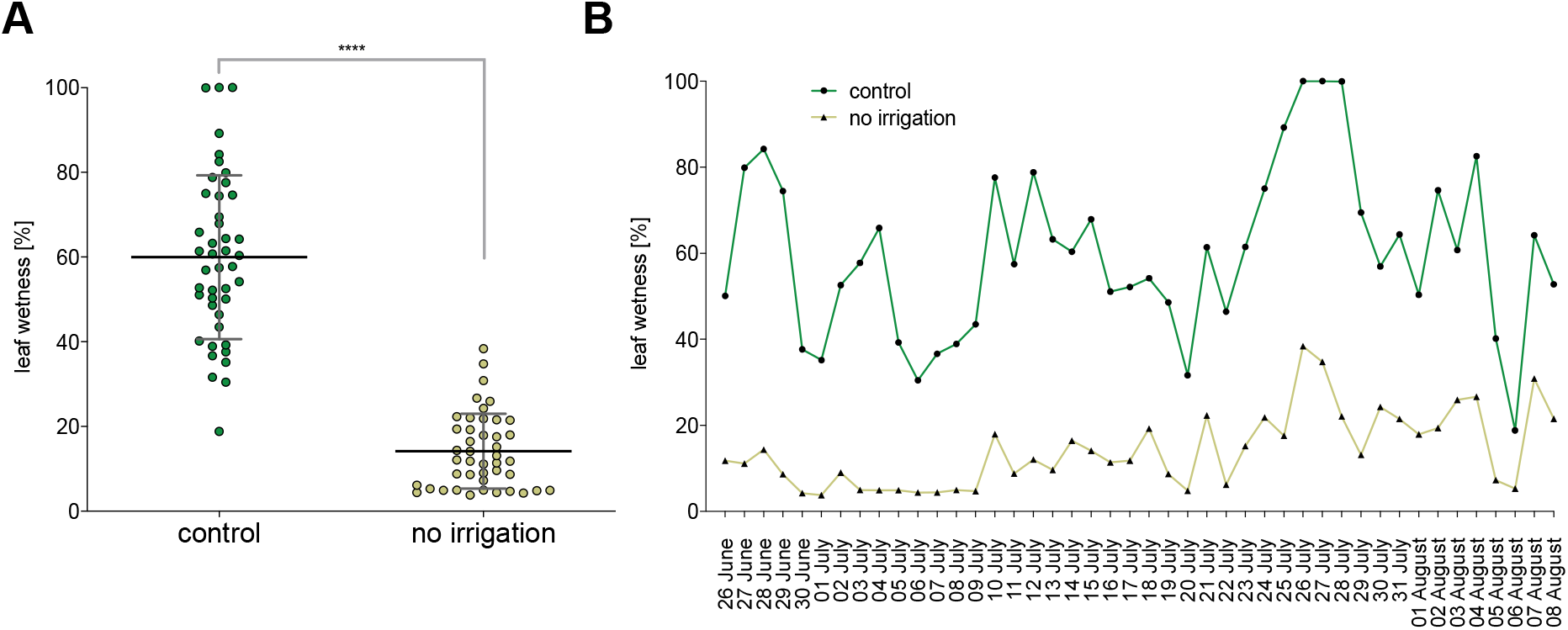
Leaf wetness of drought stressed plants compared to irrigated control monitored between 26 June and 08 August 2017 [A] and progression of leaf wetness [B] in the rainout shelter experiment. The data is presented as daily mean of leaf wetness. Error bars indicate the standard deviation of the mean. Measurements were carried out in a 10-minute interval by two leaf wetness sensors in the canopy one meter above ground. Statistical analysis: Unpaired Welch t-test (p < 0.0001).

### Monitoring of drought application in the rainout shelter

Tensiometers were used to monitor water demand and drought conditions of the barley plants in the rainout shelter. Therefore, soil moisture tension was measured three times per week in a depth of 15, 45 and 75 cm in the control and the plots with no irrigation. Exemplary data of soil moisture tension in a depth of 45 cm is presented in supplemental figure S-1. A soil moisture tension below −500 hPa was referred as drought. Drought conditions were obtained by a stop of irrigation at the begin of May. The data revealed a subsequent reduction in soil moisture in 2016 to 2018, resulting in a soil water potential below −500 hPa at end of May. During June and July, average soil moisture tension remained below −500 hPa in non-irrigated plots in all three seasons indicating a stable application of permanent and long-lasting drought conditions.

### Evaluation of Ramularia resistance under field conditions

In addition to the rainout shelter experiments, we conducted a four-year field plot trial (between 2016 and 2019) to evaluate the natural RLS infection of the same 15 spring barley genotypes. Therefore, RLS symptoms and stress-related physiological leaf spots (PLS) were evaluated as RLS leaf spot complex. The examined symptoms are shown as mean percentage of leaf area with RLS+PLS symptoms [Fig. 4a]. Symptomatic leaf areas of the upper leaves varied between 19.4 % and 33.6 % with apparent differences between the genotypes. Least symptoms appeared in the genotype Argentinische DH 168 followed by RGT Planet (21.4 %), IPZ 24727 (21.5 %) and Marnie (22.0 %). High percentage of affected leaf areas were recorded for Quench, Romilda (32.5 %) and Aischa (31.9 %). Total mean of symptomatic leaf area across the 15 genotypes was 26.3 %. Rates of RLS symptoms varied between seasons and was highest in 2016 (total mean: 45.3 %), followed by 2018 (32.3 %) and 2017 (20.8 %). RLS symptoms occurred least in 2019 (total mean: 6.9 %). In all four seasons, RLS was the dominant foliar disease over net blotch and brown rust, which both occurred at a low level [Fig. S-5].

**Figure 4.**
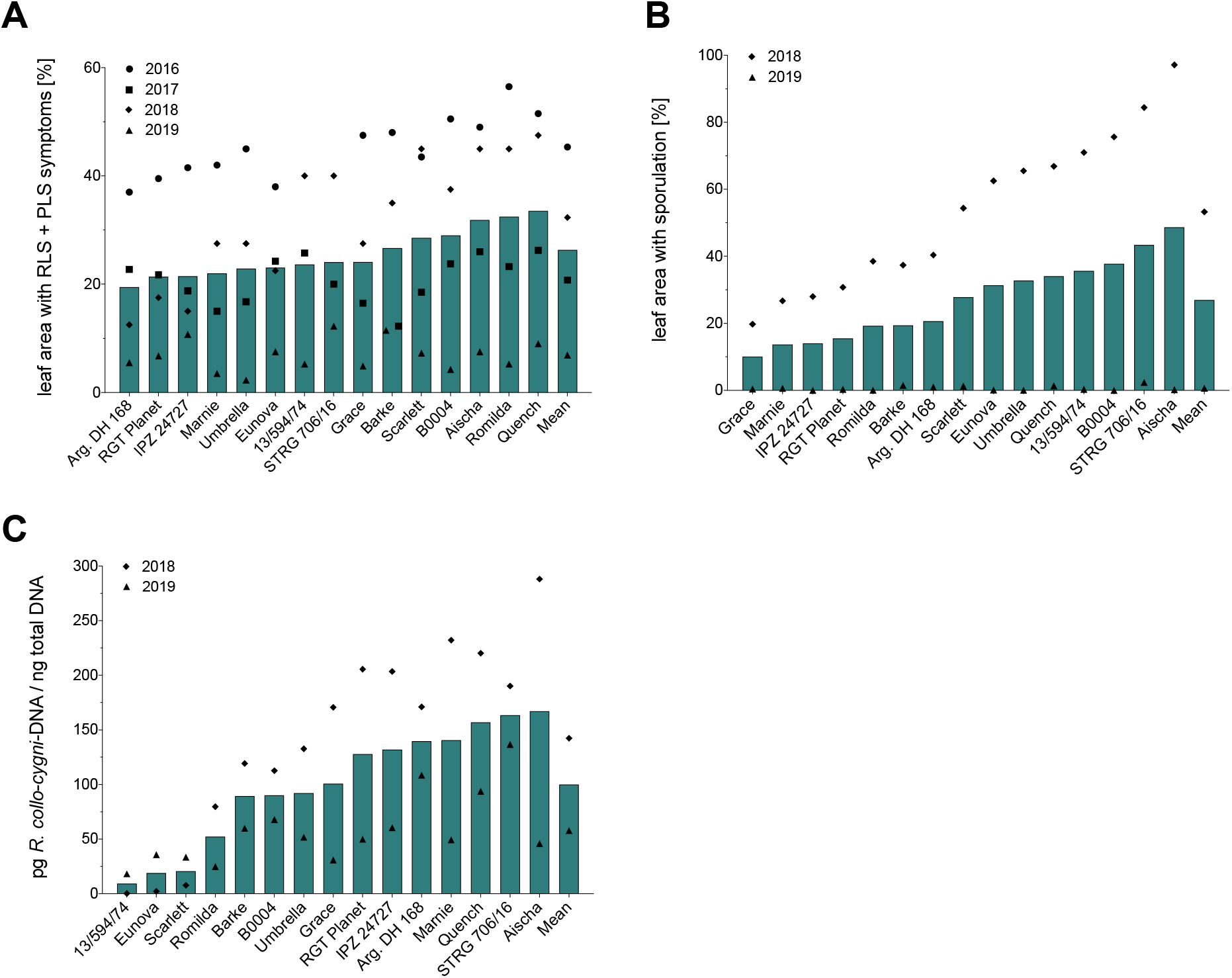
Levels of Ramularia leaf spot infestation according to assessed disease parameters in the field trials between 2016 and 2019. The bars indicate calculated mean values; symbols indicate single values. [A] represents percentage of leaf area with Ramularia and physiological leaf spots in the years 2016 to 2019. [B] indicates percentage of leaf area with sporulation in 2018 and 2019. [C] Content of Ramularia-DNA in pg Rcc DNA / ng total DNA in the upper leaves in 2018 and 2019.

In addition to visual assessments, we evaluated leaf samples for sporulation and DNA contents in the upper leaves in 2018 and 2019. The mean percentage of leaf area with visible sporulation ranged between 10.1 % in Grace and 48.7 % in Aischa. Total mean of the 15 genotypes was at 27.0 % of the leaf area showing sporulation. A high variation in total mean was observed between the two years (2018: 53.3 %; 2019: 0.6 %). [Fig. 4b]. The average content of *R. collo-cygni-DNA* in the upper leaves varied between 9.3 (13/594/74) and 167.0 (Aischa) pg/ng total DNA, which resembles a factor of 18. Besides for breeding line 13/594/74, relatively low DNA contents were also detected in Eunova (19.0 pg/ng total DNA) and Scarlett (20.6 pg/ng total DNA). The total mean of DNA contents (100.1 pg/ng total DNA) was determined from a mean of 142.4 and 57.9 pg/ng total DNA in 2018 and 2019, respectively, which indicates variation in RLS infestation between the two seasons [Fig. 4c].

Based on the collected data, we calculated a disease severity ranking (rank 1 to 15 for individual genotypes) [Fig. 5]. The lowest mean ranking (highest disease resistance) was calculated for Grace (mean rank 5.2), followed by Eunova (mean rank 5.8), Romilda (mean rank 6.0), Umbrella (mean rank 6.25) and IPZ 24727 (mean rank 6.4). High mean rankings of Quench (mean rank 12.8), breeding line STRG 706/16 (mean rank 12.6) and Aischa (mean rank 10.8) indicated susceptibility to RLS.

**Figure 5.**
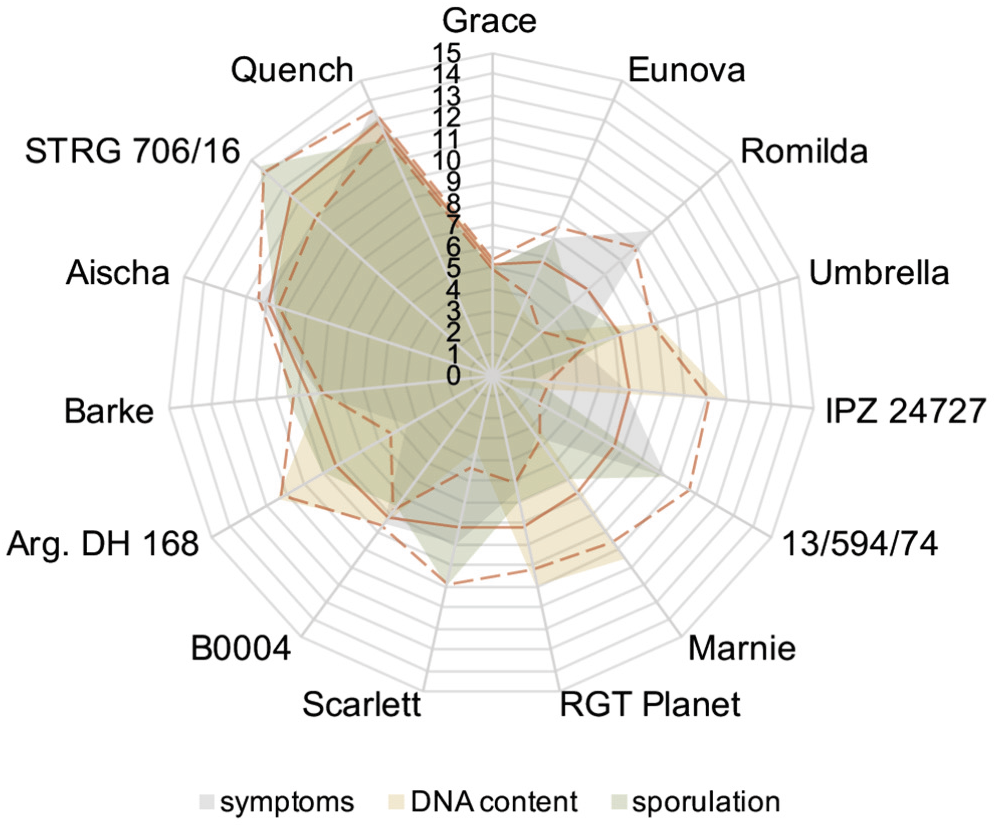
Disease severity ranking according to assessed disease parameters in the field trials between 2016 and 2019. The radar plot shows mean ranks for each disease parameter. The red lines indicate the total mean rank over individual years and disease parameters (n = 8 ranks per genotype); dashed lines indicate positive and negative standard deviation. The genotypes are sorted clockwise by total mean rank whereby low ranks indicate quantitative resistance and high ranks indicate susceptibility to Ramularia leaf spot. Individual ranks are in a range of minimum 1 and maximum 15. If two or more lines share the same rank, the same value was associated to those lines.

### Monitoring of microclimate and weather conditions in the field trails

For a better interpretation of disease severity and progression in the field, the microclimate within the canopy was monitored in the seasons 2017 to 2019 [table 2]. We recorded data during growth stages of booting and grain maturity parallel to the rainout shelter experiments. The mean temperature in the canopy near the ground was between 18.2 °C (2018) and 20.7 °C (2019). We recorded the highest mean temperature within canopy (24.4 °C) in the first week of July in 2019. Total average air humidity within the plots was 75.1 % (2017), 82.0 % (2018) and 79.8 % (2019). The mean dew point temperature was in accordance with temperature and relative air humidity.

**Table 2.** Recorded microclimate of spring barley grown in field trials in 2017, 2018 and 2019. The data was collected by data logger placed in a radiation shield on the ground within the canopy (logging interval: 1 hour). The table shows weekly mean temperature and mean relative humidity between the 2^nd^ week of June and 4^th^ week of July. The temperature of dew point was automatically determined by the data logger based on temperature and relative humidity. Measurements were conducted in time frames according to late growth stages (booting to grain maturity).

|  | temperature [°C] |  |  | relative humidity [%] |  |  | dew point [°C] |  |  |
| --- | --- | --- | --- | --- | --- | --- | --- | --- | --- |
|  | 2017 | 2018 | 2019 | 2017 | 2018 | 2019 | 2017 | 2018 | 2019 |
| 2<sup>nd</sup> week of June | 16.6 |  | 19.5 | 83.5 |  | 76.8 | 13.3 |  | 14.9 |
| 3<sup>rd</sup> week of June | 19.9 | 18.0 | 19.9 | 70.0 | 85.0 | 87.7 | 13.6 | 15.3 | 17.4 |
| 4<sup>th</sup> week of June | 22.5 | 17.2 | 21.4 | 67.7 | 79.5 | 87.1 | 15.5 | 13.4 | 18.6 |
| 1<sup>st</sup> week of July | 20.6 | 17.2 | 24.4 | 77.2 | 84.2 | 75.8 | 15.9 | 14.1 | 19.1 |
| 2<sup>nd</sup> week of July | 21.1 | 18.7 | 21.0 | 73.4 | 80.7 | 75.1 | 15.4 | 14.9 | 15.6 |
| 3<sup>rd</sup> week of July | 19.8 | 18.4 | 18.2 | 82.1 | 80.6 | 81.9 | 16.0 | 14.5 | 14.5 |
| 4<sup>th</sup> week of July | 21.6 | 19.6 | 20.5 | 71.0 | 82.8 | 75.6 | 15.1 | 15.9 | 15.4 |
| average | 20.4 | 18.2 | 20.7 | 75.1 | 82.0 | 79.8 | 15.1 | 14.7 | 16.4 |

We further collected weather data from a nearby weather station for the years 2016-2019. For details of the seasonal weather conditions in respect to average monthly temperature, average monthly precipitation, average monthly leaf wetness and average monthly radiation, we refer to supplemental tables S3-S6.

### Correlation of leaf senescence and RLS disease parameter

In parallel to RLS symptoms, we had recorded leaf senescence in the rainout shelter and in the field. To assess a relation between leaf senescence and further ripening with RLS disease parameters, the obtained data of senescent leaf area and RLS disease was used to calculate Pearson’s correlations for the rainout shelter experiment and field trials [Fig. S-4]. A positive significant correlation occurred between senescent leaf area and Ramularia-DNA in the irrigated control of the rainout shelter. In the field trials, leaf senescence was significantly negatively correlated with RLS symptoms and sporulation but not with fungal DNA [Figs. S-4c, S-4f].

### Evaluation of genotype stability for yield parameters

In order to determine the stability of genotypes according to yield parameters across environments a stability index (yield parameter stability index, YPSI) was calculated, based on data of yield, fraction of kernels with a size > 2.5 mm, fraction of kernels with a size > 2.8 mm and thousand kernel weight (TKW) [Fig. 6]. The lower the YPSI is, the more stable is the performance of a certain genotype for the considered yield parameters over several environments. The cultivar Aischa and breeding line B0004 showed the highest stability (lowest YPSI) across the determined yield parameters (YPSI: 172; YPSI: 181), whereas IPZ 24727 (YPSI 363) and Barke (YPSI 351) were least stable. Total mean of YPSI over all 15 tested genotypes was 637.5. Marnie and Grace scored highest stability in thousand kernel weight (YPSI: 36; GSI: 39), Romilda (YPSI: 29) and Aischa (YPSI: 30) for yield.

**Figure 6:**
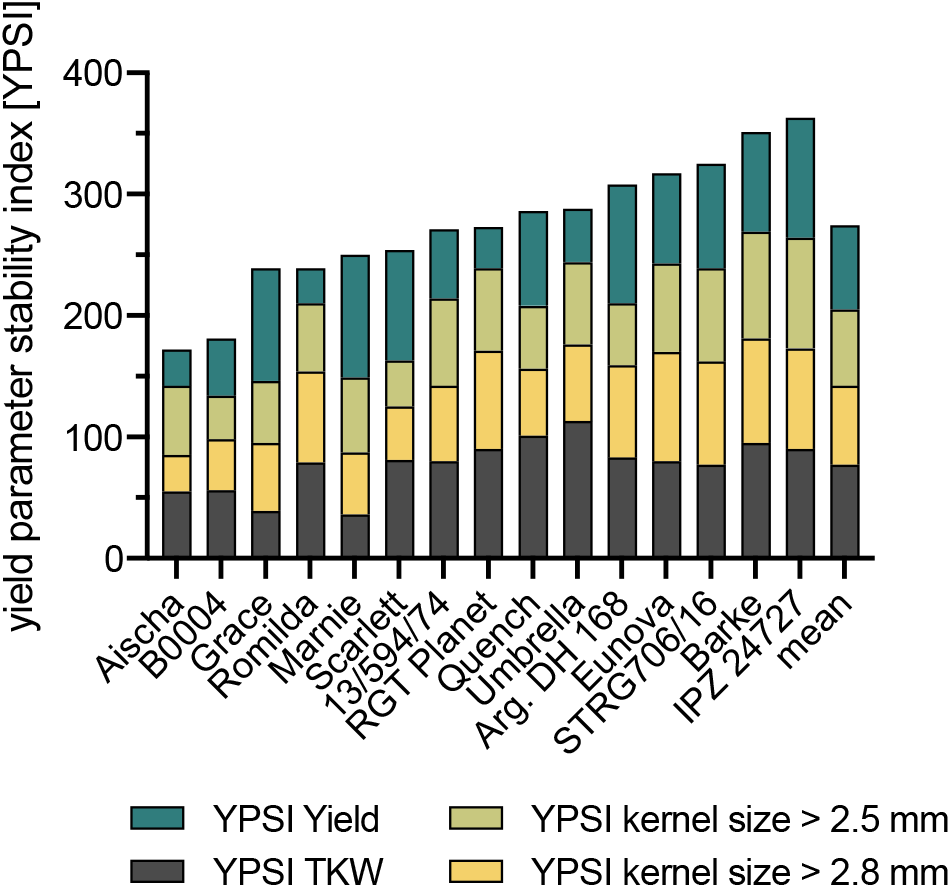
Yield parameter stability index [YPSI] calculated for the assessed yield parameter: yield, thousand kernel weight (TKW), kernel size fraction > 2.5 mm and kernel size fraction > 2.8 mm. The yield parameters were determined at nine locations including the rainout shelter, field trial (Freising) and six additional field locations in 2018. YPSI of each genotype was summed up for all yield parameters. The genotype with the smallest YPSI was considered to be the best across yield parameters and different field locations.

## Discussion

Understanding the impact of environmental conditions on epidemiology of diseases and on resistance of crop plants is crucial for development of multiple stress resistant genotypes under aspects of climate change. The aim of this study was to screen an assortment of spring barley genotypes for basal resistance against *R. collo-cygni* under natural, irrigated and drought conditions in the field. Data show a cross-talk between drought and the RLS disease and strong differentiation of quantitative RLS resistance in spring barley genotypes. This may ease choice of cultivars by farmers and support efforts to breed for RLS resistance.

The rainout shelter experiment provided data about genotype performance and RLS resistance under both, drought conditions and irrigation over three consecutive field seasons. The obtained data revealed a clear differentiation in disease severity between the most resistant and most susceptible genotypes, especially in the irrigated control. A few genotypes showed a reproducible RLS resistance over three consecutive years. Furthermore, the tested genotypes represent a diverse set of largely modern barley cultivars commonly grown in Europe and current breeding material from German barley breeders. Hence, data identify available cultivars directly recommendable for practical farming, although, of course, even more RLS-resistant genotypes might be available that have not been tested here.

The genotypes IPZ 24727, Grace, Umbrella and Eunova exhibited little symptoms, sporulation and low fungal DNA contents in the upper leaves both in the irrigated control and under drought stress. This indicates a robust and drought stress-independent level of quantitative resistance against RLS disease. Apart from these cultivars, the effect of the environment on other genotypes tended to be intense. High year-to-year differences in leaf spotting, sporulation and DNA contents [Fig. 1b, 1d, 1f] suggest that uncontrolled weather parameters and potentially spore flight influenced disease severity particularly in more susceptible genotypes. In particular, disease severity was highest in 2016, where weather conditions were different from the following years (e.g. cooler weather conditions [Table S-3]). To balance the high variation in disease severity between the considered years, we calculated ranks for each disease parameter and a mean ranking for RLS resistance for each genotype [Fig. 2]. According to our data, the ranking revealed a reproducible RLS resistance (e.g. breeding line IPZ 24727, cultivar Umbrella), or susceptibility (e.g. cultivar Scarlett), respectively, of several genotypes under irrigation and drought conditions. Hence, the ranking is an applicable approach to assess resistance obtained from different diagnostic tools for the selection of genotypes over several seasons, despite high year-to-year variation in obtained data. Furthermore, some genotypes referred as susceptible under irrigation were less affected by RLS under drought stress, e.g. cultivar Barke and Quench, which is represented by appropriate rankings [Fig. 1, 2]. It should be noted, that the quantitative RLS resistance and susceptibility reported here was stable over several seasons for several genotypes, but we cannot predict whether this would generally occur at other locations (compare also field and rainout shelter ranks of certain lines such as Scarlett). The strong impact of weather and possibly microclimate conditions on the RLS disease may also lead to unexpected environment-genotype interactions under conditions not tested here. Hence, further disease monitoring of a diverse panel of genotypes and at diverse locations seems recommendable for breeders and consultants.

Dry conditions in the rainout shelter were unfavourable for the development of RLS. Possibly, spore germination, infection through stomata and proliferation of *R. collo-cygni* were suppressed. Duration of stomata closure as a response to drought stress might have an impact on the rate of secondary infections by Ramularia conidia, which was shown to be important for pathogenesis (Stabentheimer et al., 2009). We further speculate that under prolonged drought conditions, the accelerated senescence of the host plant could have limited the time for successful pathogenesis of RLS. Under the applied extreme permanent drought conditions, plant development until full ripening occurred faster, which may have interfered with pathogen colonization of the infected leaf tissue. This may occur despite the fact that plant stress can also support the disease, potentially by weakening the host (Schützendübel et al., 2008). For the *R. collo-cygni* - barley pathosystem, it is stated that host plant ontogenesis is directly linked to disease progression (Schützendübel et al., 2008; Havis et al., 2015), which is contrasting to adult plant resistance that is e.g. observed in selected cereal genotypes to powdery mildew and other diseases (Develey-Rivière & Galiana, 2007). It seems possible, that the pathogen life style switch, from endophytic to necrotrophic, could not occur under heavy drought conditions. Drought stress may have modified plant physiology, so that the fungus could hardly benefit from quickly senescing host tissue, though it has a necrotrophic growth potential and is even able to induce accelerated leaf senescence (Sjokvist et al., 2019). We further speculate that premature senescence under drought conditions inhibits the formation of typical necrotic spots, because total leaf necrosis as a response to drought occurred before the pathogen could induce local cell chlorosis and subsequent necrosis in the leaf tissue. Severe drought thus caused relatively RLS-symptomless but senescent leaves, even though the pathogen was present at a low level. As a response to drought, an accelerated remobilization of nutrients from leaves to generative organs (Zimmermann & Zentgraf, 2005) could have withdrawn resources from infected leaf tissue and thereby limited access of *R. collo-cygni* to nutrient supply. Furthermore, responses to drought and heat at a cellular level, e.g. osmotic adjustment (Blum, 2017), plant stress altered hormone signalling, formation of reactive oxygen species (Chaves et al., 2003; Zimmermann & Zentgraf, 2005) or accumulation of antioxidants (Sallam et al., 2019) could probably impair development of *R. collo-cygni* resulting in less disease progression.

In context of RLS, it is not fully clear from literature, whether barley physiologically exhibits an age-related susceptibility to RLS independent of additional exposure to stress, e.g. by a metabolic switch to the generative development. Alternatively, the pathogen could exploit a weakened host that faces multiple stresses late in the season and hence exerts host stress-dependent virulence. There is considerable evidence that age-related physiological alterations including the breakdown of the antioxidative system within host tissue increases the imbalance between symptomless pathogen development and the active plant defence system (Heiser et al., 2003; Miethbauer et al., 2006; Schützendübel et al., 2008), resulting in a typical late but rapid outbreak of RLS disease. That most probably holds true under normal field conditions, but is strongly modulated under heavy drought stress with accelerated ripening and plant senescence. Less spotted leaf area and fungal DNA contents in leaves of drought stressed plants provide evidence for a suppression of RLS disease under very unfavourable environmental conditions supporting accelerated plant death [Fig. 1, S-2, S-4]. It remains unclear, why physiological leaf spots (PLS) were also considerably less frequent on drought stressed plants. We found a general reduction in leaf spotting including PLS on all drought stressed genotypes, even though plants normally show a higher expression of PLS caused by stress related oxidative disorder in leaf tissue (Wu & von Tiedemann, 2002). On the other hand, the tested genotypes were partially preselected for their adaptation to environmental stress, including less leaf spotting in general, which could explain lower expression of leaf spots than expected from an unbiased assortment. Furthermore, our data supports previously reported genotype-dependent expression of leaf spots (Wu und von Tiedemann, 2004). Nevertheless, it is still not clear, whether an increase of plant stress enhances RLS or is a result of RLS disease and hence is maybe involved in pathogenesis (McGrann et al., 2015). Furthermore, a survey of McGrann et al. (2015) demonstrated that a stress-responsive *NAC1* transcription factor involved in drought tolerance and plant development is affecting susceptibility to RLS. *NAC1* over-expression lines, which are more drought tolerant, simultaneously exhibited less leaf senescence, less RLS symptoms and fungal DNA in barley leaves, which further connects abiotic stress resistance and RLS resistance. By contrast, our data did not reveal a fully conclusive picture of correlation of genotype-dependent leaf senescence or green leaf area in upper leaves during ripening and RLS disease parameters in the rainout shelter when compared to the field [Figs. S-2, S-4]. At the moment we can only speculate that on the one hand, more fungal biomass in upper leaves has contributed to accelerated leaf senescence in the rainout shelter. On the other hand, early senescence in the field might have limited symptom formation and fungal sporulation [Fig. S-4c, S-4f]. The future question arises whether there is a trade off between resistance to RLS and late plant ripening. The longer leaves can maintain their function to provide assimilates and do not become senescent, the more yield is potentially produced (Gregersen et al., 2008). Prolonged juvenility might support resilience to environmental stress situations, but could also be exploited by the partially endophytic fungus to complete its life cycle. Our data on stability of yield parameters across several environments (including the rainout shelter and our field trials) [Fig. 6] suggest that there are a few genotypes that are comparably resistant to RLS and show a high yield stability (e.g. Grace, Romilda). However, the quite RLS resistant genotype IPZ 24727 revealed low stability of yield parameters over diverse field environments. Additionally, Scarlett and B0004 are comparably susceptible to RLS but still show high yield. Taken together, the question arises whether there might be a trade off between genotype-dependent ripening, yield performance and disease resistance, which should be taken into account by research and plant breeders. In this context, disease tolerance, instead of resistance, could become a potential breeding aim.

Rainfalls, air humidity, and leaf wetness duration are generally important for the dispersal, latency period, infestation and proliferation of several plant diseases caused by barley pathogens like *Pyrenophora teres* (Shaw, 1986). Several studies indicated that moisture seems to have a high impact on epidemics of RLS, too (Formayer et al, 2002; Brown et al., 2014; Havis et al., 2015). According to observations of Formayer et al. (2002), artificial prevention of leaf wetness strongly decreased symptom formation under semi-controlled conditions in the field, whereas leaf wetness by dew was sufficient for disease development. Brown et al. (2014) found a positive relationship between area under RLS disease progress curve and leaf wetness at different locations and over several years. The results are in line with our own findings in the rainout shelter experiment, where plants were not irrigated from the booting stage till harvest and further experienced increasing drought conditions [Fig. S-1]. Hence, leaf wetness exclusively originated from air humidity during the night and at low level [Fig. 3; Tab. 1]. Microclimatic measurements revealed that relative air humidity within canopy of drought stressed plants was continuously decreased, while temperature was increased when compared to irrigated plots [Table 1]. Consequently, leaf wetness was strongly reduced within the dry canopy. Simultaneously, conidiophores and conidia were hardly present on senescent leaves of drought-stressed barley. This implies that the extent and duration of remaining leaf wetness was likely insufficient for the pathogen to complete its life cycle. The same could hold true for spore germination and early colonization from air borne inoculum. Results by Havis et al. (2012) showed, that an increase of leaf wetness was followed by a massive release of spores into the air. By contrast, Schützendübel et al. (2008) observed two peaks of airborne spores in one winter barley trial during a dry period late in the season and independently from rainfall events. Hence, spore formation may primarily depend on sufficient air humidity and dew, but rain events may act as an enhancing factor. Despite the increased temperature in drought stressed canopy, a direct effect of temperature on RLS is not obvious in our data, but should have further shortened duration of leaf wetness. Taken together, our data supports the importance of moisture for sporulation of *R. collo-cygni* and shows a strong suppression of RLS development under long-lasting drought conditions.

In addition to stress experiments in the rainout shelter, the spring barley genotypes were pathophenotyped in parallel field trials over four years (2016-2019). In general, disease incidence varied between seasons and revealed a high level of symptoms, sporulation and fungal DNA contents in upper leaves, thus indicating strong natural disease pressure at the location. At late growth stage 83-85, RLS and PLS formed the predominant foliar disease ahead of net blotch and brown rust [Fig. S-5], but differences in symptomatic leaf areas between genotypes were less noticeable when compared to the rainout shelter [Fig. 1]. The assessment of fungal sporulation and determination of Ramularia DNA revealed stronger differences in resistance to *R. collo-cygni* between the tested spring barley genotypes compared to the rainout shelter experiment [Fig. 1, 4]. However, it is difficult to assess field resistance to RLS from single factors, e.g. symptom monitoring without determination of fungal DNA contents. Hence, the applied diagnostic tools complement each other and increased robustness of field data. Based on the three recorded disease parameters, a calculation of mean ranks revealed comparable disease resistance of several genotypes in both open field and rainout shelter experiments. Thus, scoring of RLS parameters under field conditions confirmed resistance of several genotypes that was observed under semi-controlled conditions in the rainout shelter (e.g. IPZ 24727, Grace, Eunova and Umbrella). Furthermore, measurements of meteorological factors including temperature, precipitation and relative air humidity on site and by a nearby weather station revealed mainly similar conditions in the field trials compared to the irrigated control in the rainout shelter experiment. By contrast, only leaf wetness was higher in the rainout shelter than in the field during the recorded period [Fig. 3, Tab. S-4], probably due to weekly irrigation of 20 mm. By comparing disease parameters with climatic conditions of individual field seasons, there was no clear cross-talk between warm and dry conditions and RLS severity. Highest RLS severity across all disease parameter was observed in 2016, where temperatures were cooler than in the following seasons. RLS severity was indeed lower in the warmer and dryer years 2017 and 2019, but not in 2018. In fact, very hot periods after barley flowering stage in June and July 2018 were intermitted by single rain events, possibly allowing for strong disease progression. Hence, single rain events could likely provoke sufficiently moist conditions. Furthermore, high air humidity during the night enhance leaf wetness by dew formation, which we could observe in particular in our field trials in 2018. However, climate change will likely cause more drought periods in central Europe and Germany (Zebisch et al., 2005) and thus will further affect plant diseases. For the case of RLS, our findings suggest that dry and hot conditions could suppress RLS disease in years with very long-lasting lack of rain. However, there are also reasons to assume that single rain events in dry summers such as 2018 are sufficient to enhance RLS disease of pre-stressed plants.

To meet the challenge of better understanding the complex interaction of host genotypes, environment and disease progression, this study provides a first insight into multiple stress resistance of barley under aspects of a changing climate. Further experiments from controlled and complex environments may help future prediction of RLS disease dependent on the cultivar and seasonal weather conditions. Our data are relevant for crop production and future breeding of robust and resilient crop plants. In particular, the current lack of effective fungicides to control RLS reinforces the importance of resistance breeding and the choice of quantitative RLS resistant barley cultivars. Indeed, we identified a set of spring barley genotypes that were quite resistant to RLS under three different field environments in three years of investigation.

## Conclusions

Since the 1980s, *Ramularia collo-cygni* became a major pathogen in barley production around the world. The possible connection between disease progression, environmental conditions and plant maturity is still not fully understood. Genetic resistance of barley to RLS is limited and hardly exploited. The present study describes suppressive effects of strong drought conditions on RLS severity of a preselected panel of spring barley genotypes. We identified genotypes with a reliable resistance to RLS disease in a multi-annual study under controlled drought, irrigated and natural conditions in the field. The results support that RLS depends on moisture and is suppressed under persistent drought conditions that rarely occur under open field conditions. Parallel field trials during very warm and dry seasons suggest that the formation of dew and single rain events still suffice for considerable RLS disease outbreaks. Moreover, our investigations contribute to a better general understanding of interactions between environmental factors and the epidemics of RLS. The study further validated the value of complementary diagnostic tools for phenotyping of barley RLS resistance.

## Acknowledgements and funding

This project was financially supported by the Bavarian State Ministry of the Environment and Consumer Protection in frame of the Project network BayKlimaFit (www.bayklimafit.de); subproject 10: TGC01GCUFuE69781.

We thank the Ackermann Saatzucht GmbH & Co. KG (Irlbach, Germany), Saatzucht Josef Breun GmbH & Co. KG (Herzogenaurach, Germany) and Saatzucht Streng-Engelen GmbH & Co. KG (Uffenheim, Germany) for good corporation and providing and propagating seed material. We are greatful to Dr. Anja Hanemann (Saatzucht Breun GmbH & Co. KG), Christian Schuy (Friedrich-Alexander University Erlangen-Nürnberg) and Prof. Dr. Lars Voll (Philipps University of Marburg) for their comprehensive support in selecting barley genotypes.

The authors thank the field technician team of the greenhouse and laboratory center Dürnast (Technische Universität München) and the field technician team of the Bavarian State Institute of Crop Science and Plant Breeding in Freising for maintenance of field experiments. We also want to thank Carolin Hutter, Alice Retter and Bianca Eibl for technical support.

## Supplements

**Figure S-1.**
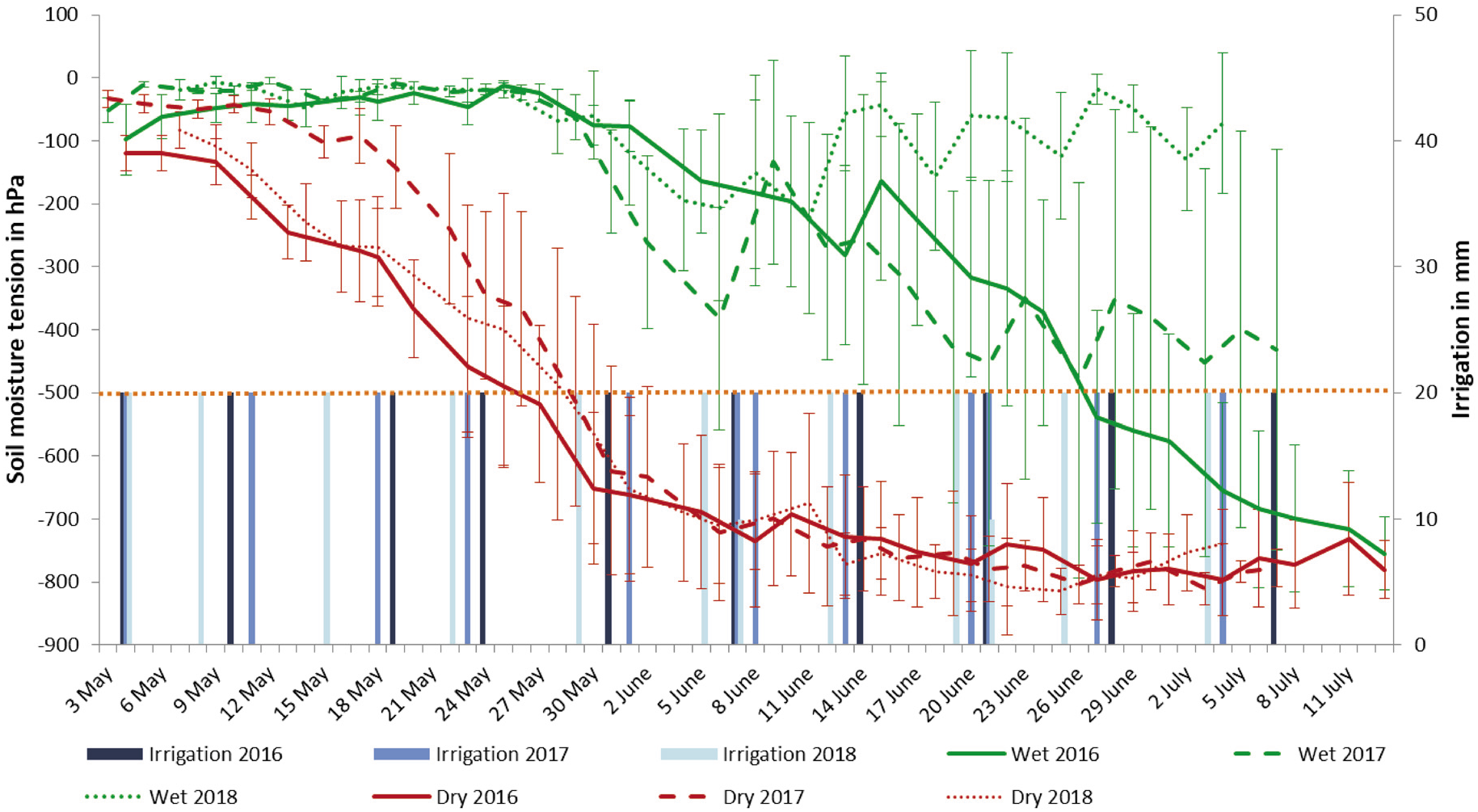
Soil moisture tension in a depth of 45 cm in the rainout shelter experiment in a period between May and July in 2016 to 2018. Soil moisture is represented in hPa. Error bars show the standard error of the mean. Drought condition was set as soil moisture tension below −500 hPa. The weekly irrigation in the control plots was 20 mm and is represented as blue bars.

**Figure S-2:**
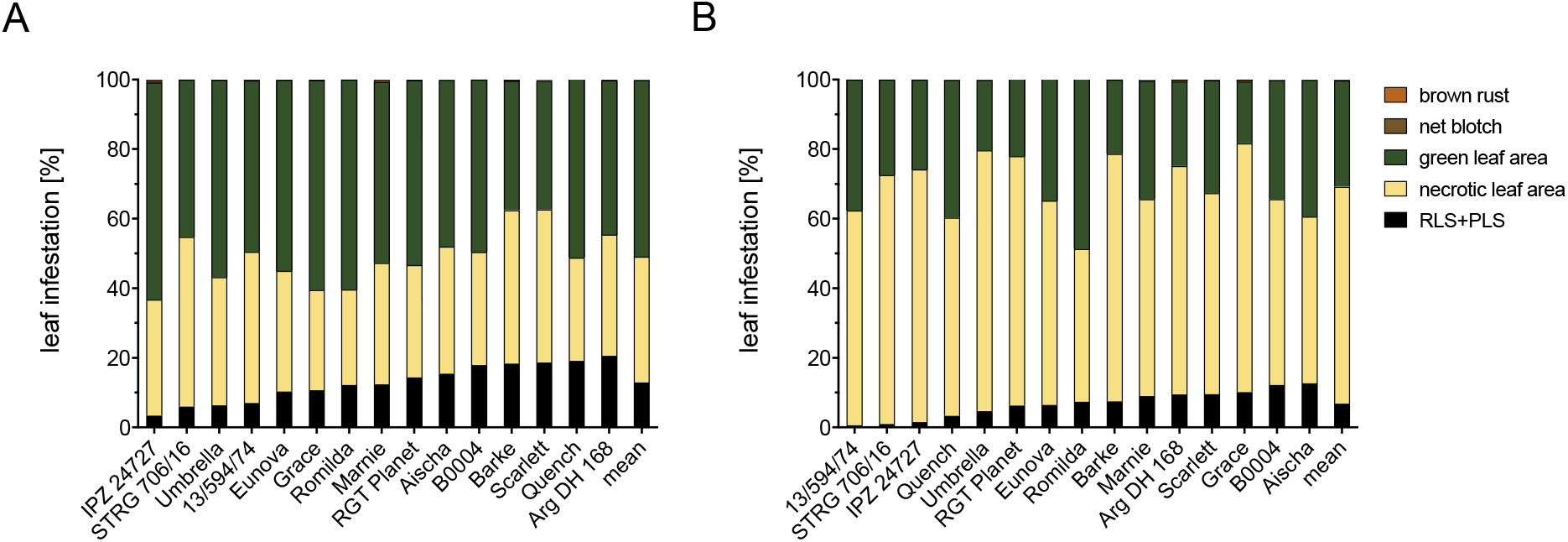
Mean proportions of green leaf area, senescent leaf area and infestation of recorded foliar diseases (brown rust, Ramularia and physiological leaf spot, net blotch) on upper leaves (flag leaf, flag leaf −1) of 15 spring barley genotypes grown under controlled conditions [A] or under drought conditions [B] in the rainout shelter between 2016 and 2018. Visual assessment was conducted at growth stage 83 – 85.

**Figure S-3.**
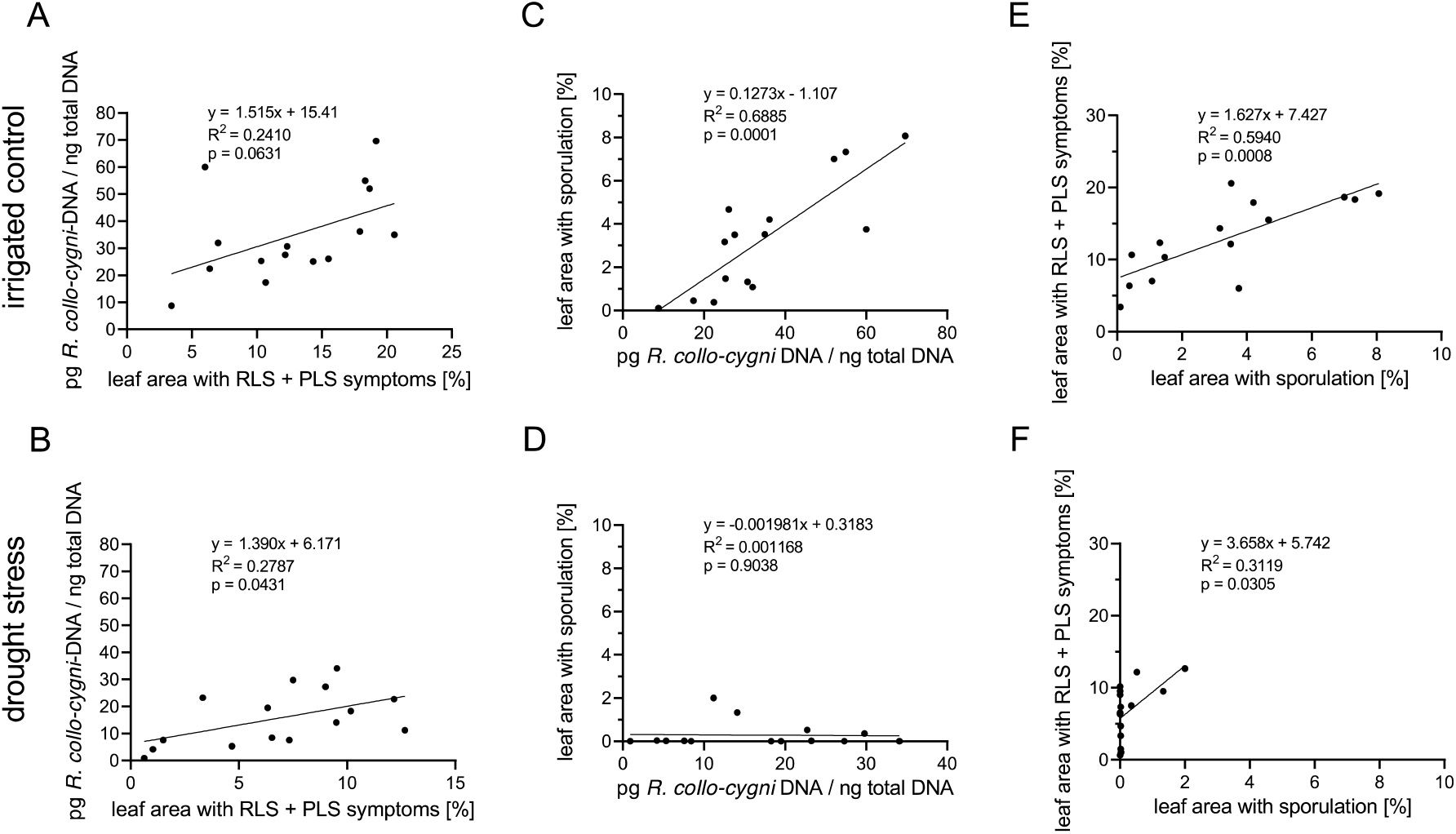
Relations between assessed Ramularia leaf spot disease parameters in the rainout shelter between 2016 and 2018. Data points represent calculated means of 15 spring barley genotypes. RLS and PLS symptoms and Ramularia-sporulation are presented as percentage of leaf area, respectively. Visual assessment of upper leaves was conducted at growth stage 83 – 85. Statistical analysis: linear regression; p value indicates significance of the slope from zero; alpha = 0.05.

**Figure S-4.**
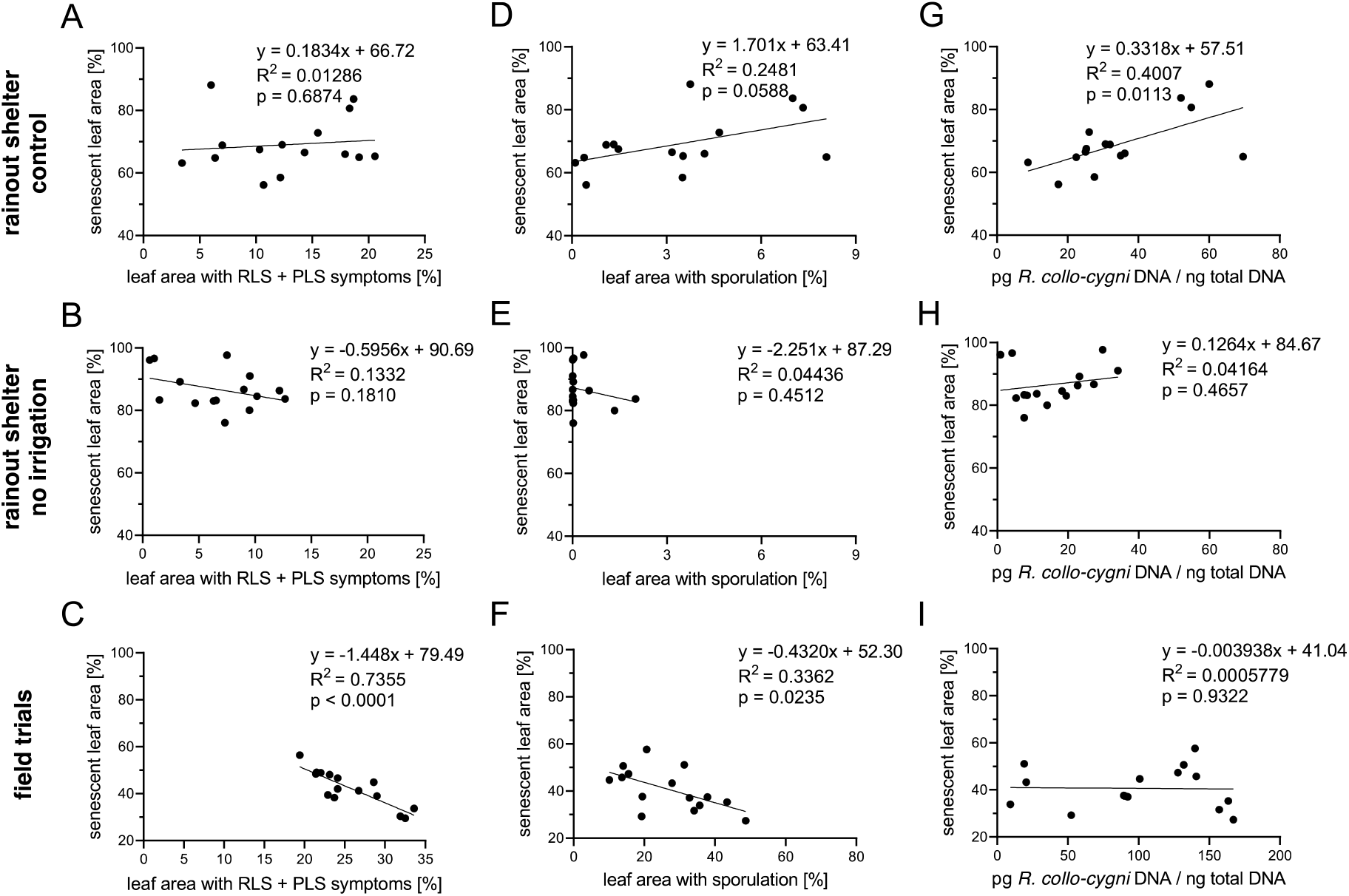
Relations between assessed leaf senescence of flag leaf and flag leaf minus one (F-1) and Ramularia leaf spot disease in the rainout shelter between 2016 and 2018 and in the field trials between 2016 and 2019. Data points represent calculated means of 15 spring barley genotypes. Leaf senescence, symptoms of RLS and PLS and Ramularia-sporulation are presented as percentage of leaf area, respectively. Visual assessment of upper leaves was conducted at growth stage 83 – 85. Statistical analysis: linear regression; p value indicates significance of the slope from zero; alpha = 0.05.

**Figure S-5.**
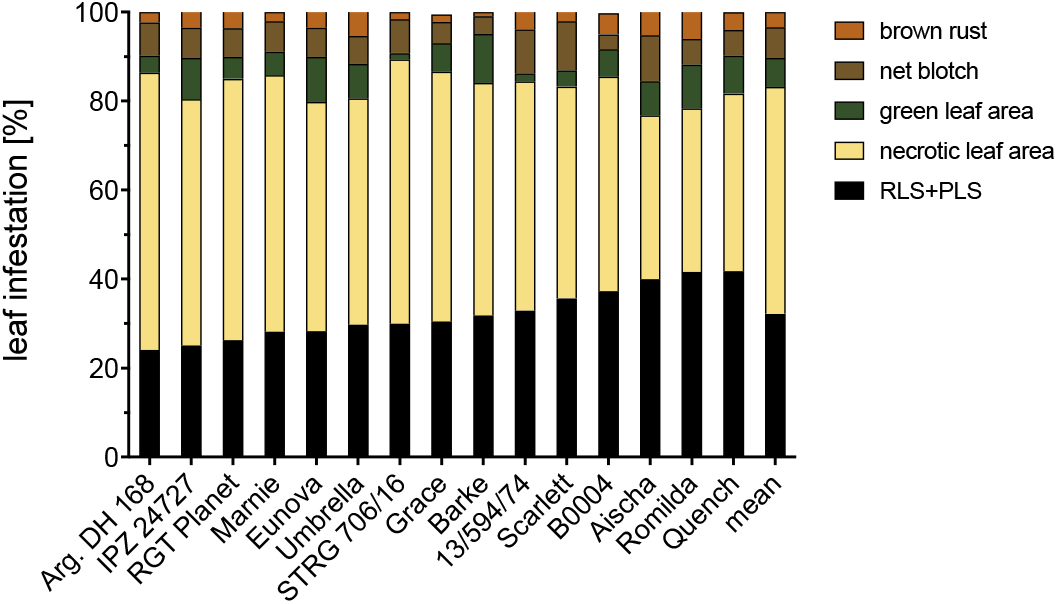
Mean proportions of green leaf area, senescent leaf area and infestation of recorded foliar diseases (brown rust, Ramularia and physiological leaf spot, net blotch) on upper leaves (flag leaf, flag leaf −1) of 15 spring barley genotypes grown in the field trials between 2016 and 2019. Visual assessment was conducted at growth stage 83 – 85.

**Table S-1.**
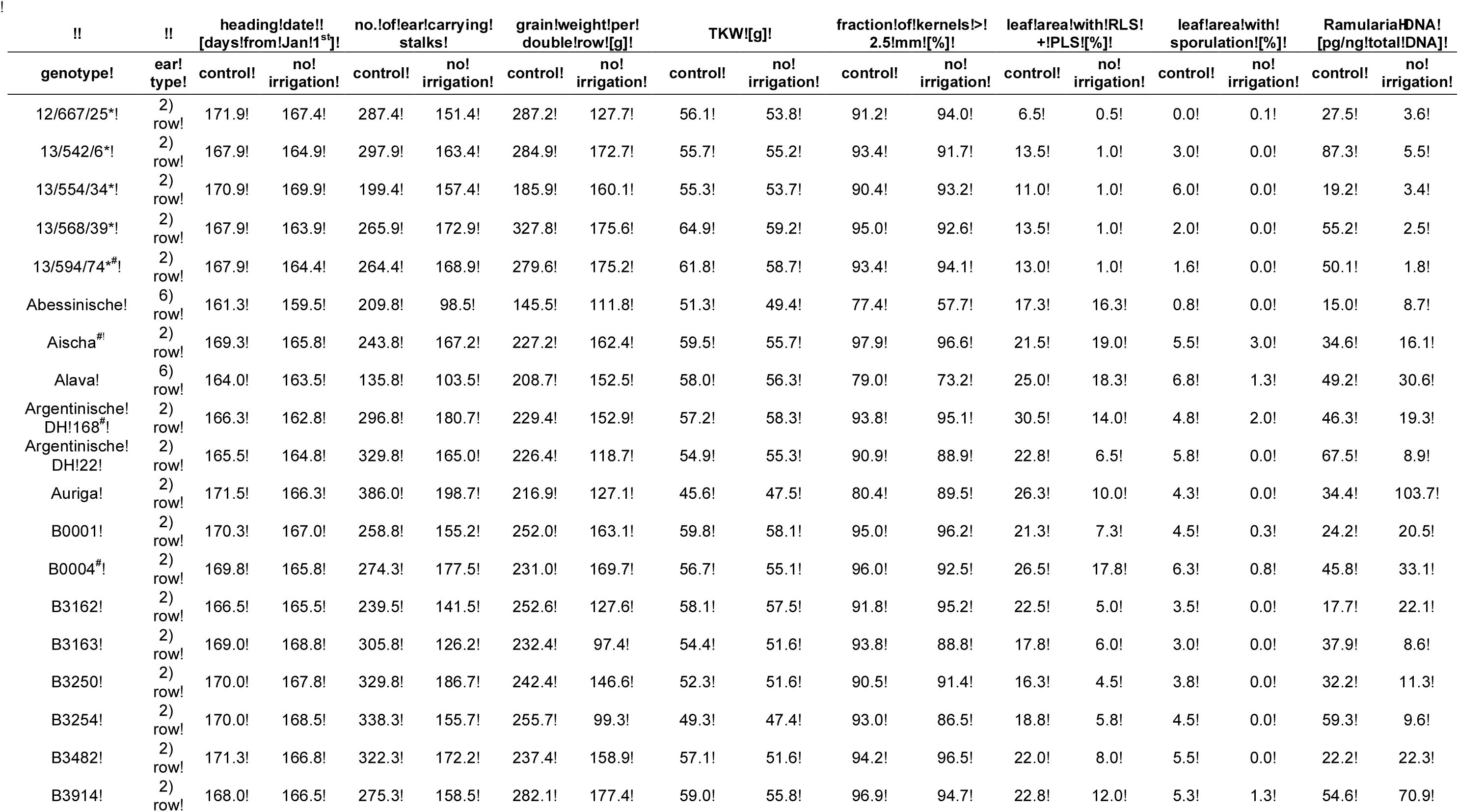
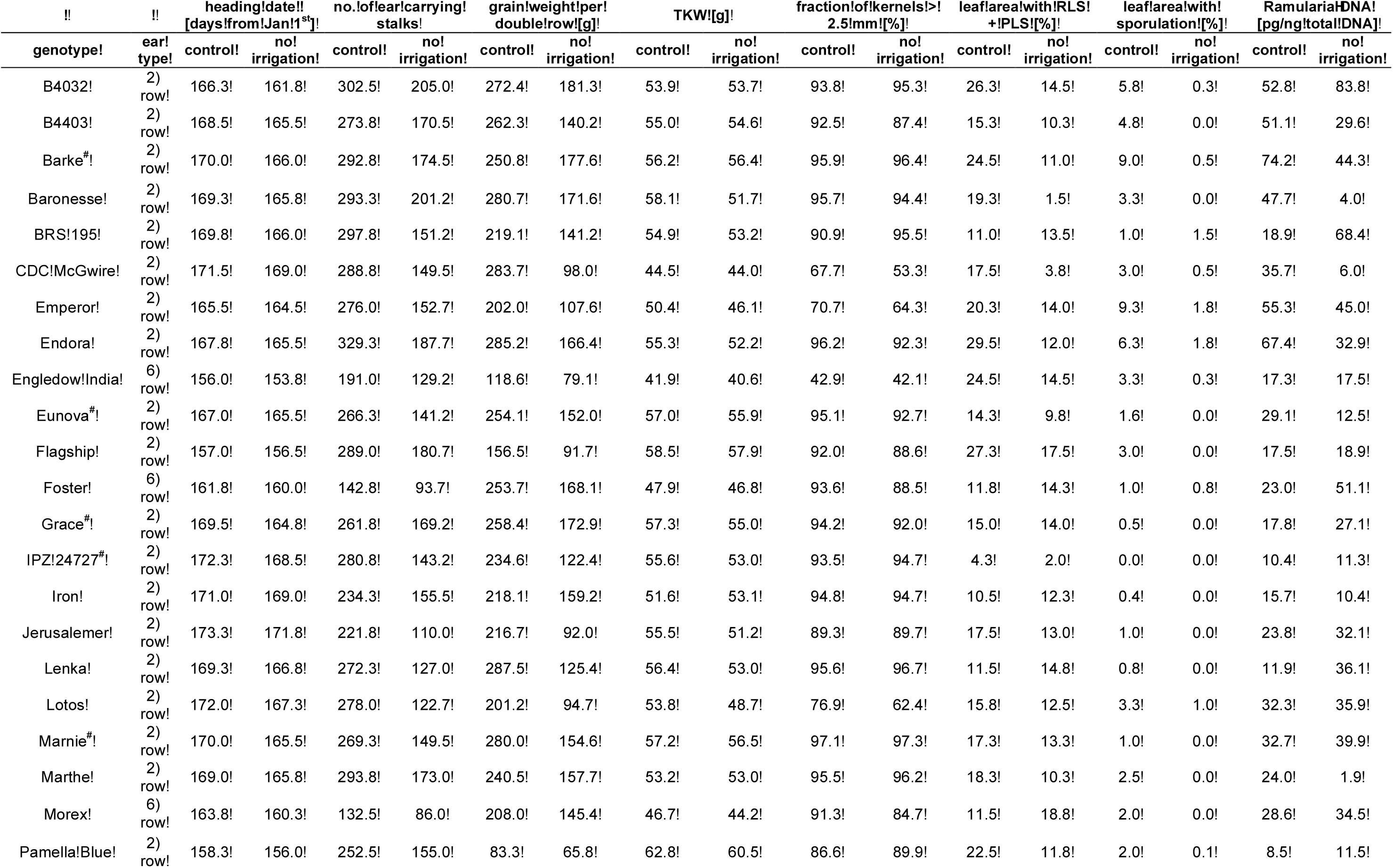
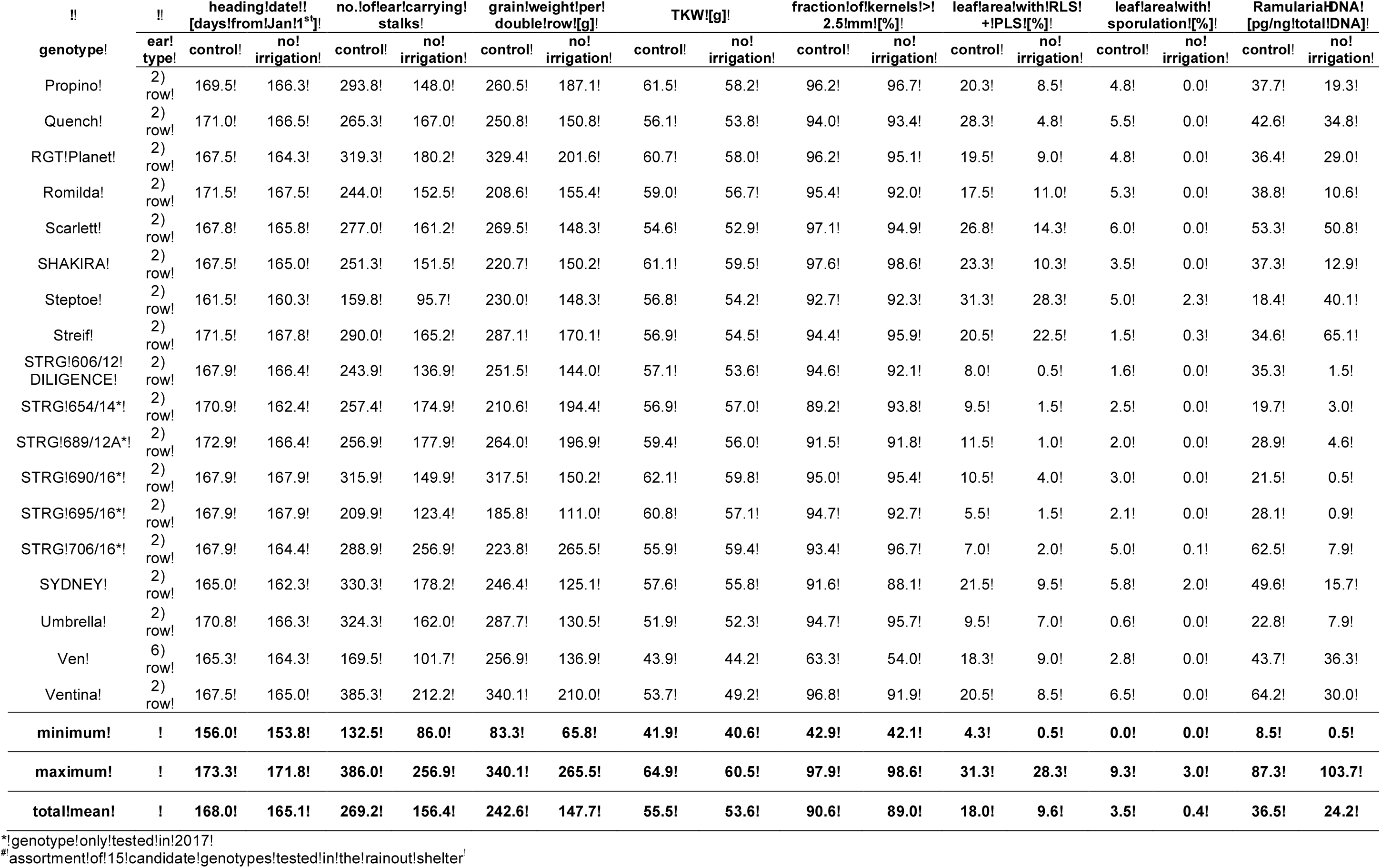
Agronomic data and RLS disease infestation of 59 spring barley genotypes tested in double rows (micro plots) under irrigation (control) and no irrigation in the rainout shelter in 2016 and 2017. The table demonstrates the mean date of heading [days from Jan 1^st^], mean number of ear carrying stalks, average of total grain weight per double row [g], mean thousand kernel weight (TKW) [g], the mean of the fraction of kernels with a size > 2.5 mm [%], mean leaf area with RLS and PLS symptoms [%], mean leaf area with sporulation [%] and mean Ramularia-DNA contents [pg/ng total DNA] of flag leaf (F) and flag leaf minus one (F-1).

**Table S-2.** Dates of sowing, growth stages, harvest and number of tested genotypes in field trials and in the rainout shelter experiment in seasons between 2016 and 2019.

|  | field trial |  |  |  | rainout shelter |  |  |
| --- | --- | --- | --- | --- | --- | --- | --- |
|  | 2016 | 2017 | 2018 | 2019 | 2016 | 2017 | 2018 |
|  | microplots | microplots | field plots | field plots | microplots | microplots | field plots |
| number of tested genotypes | 59 | 59 | 59 | 15 | 59 | 59 | 15 |
| 3rd week of March |  | sowing |  | sowing |  |  |  |
| 4th week of March |  |  |  |  |  | sowing |  |
| 1st week of April |  |  |  |  | sowing |  |  |
| 2nd week of April |  |  | sowing |  |  |  | sowing |
| 3rd week of April | sowing |  |  |  |  |  |  |
| 4th week of April |  |  |  |  |  |  |  |
| 1st week of May |  |  |  | GS 49 |  |  |  |
| 2nd week of May |  |  |  |  |  |  |  |
| 3rd week of May |  |  |  |  |  |  |  |
| 4th week of May |  |  |  |  |  |  |  |
| 1st week of June |  | GS 32 | GS 55-59 |  | GS 51 | GS 51 |  |
| 2nd week of June | GS 32 |  |  | GS 51-55 |  |  |  |
| 3rd week of June | GS 37 |  | GS 73-83 | GS 63-67 |  |  |  |
| 4th week of June |  |  |  |  |  |  | GS 73-83 |
| 1st week of July | GS 49-71 |  | GS 85-87 | GS 85-87 |  |  |  |
| 2nd week of July |  |  |  |  |  |  | GS 85-90 |
| 3rd week of July |  |  |  |  | harvest | GS 90 |  |
| 4th week of July |  |  |  |  |  | harvest |  |
| 1st week of August |  |  |  |  |  |  |  |
| 2nd week of August |  | GS 90 |  |  |  |  |  |
| 3rd week of August | GS 90 |  |  |  |  |  |  |

**Table S-3.** Recorded mean temperature in two meters above ground during field seasons between 2016 and 2019 and calculated means for each month between March and July. Mean temperatures between 1961 and 1990 are listed as baseline. The data was collected by a weather station at the location in Freising. The data was accessed from agro-meteorology web portal of the Bavarian State Institute for Agriculture (Agrarmeteorologie Bayern, 2020).

|  | average temperature [°C] |  |  |  |  |  |
| --- | --- | --- | --- | --- | --- | --- |
|  | 2016 | 2017 | 2018 | 2019 | average (2016 - 2019) | average (1961-1990) |
| March | 3.9 | 6.9 | 3.0 | 6.3 | 5.0 | 3.0 |
| April | 8.2 | 7.4 | 13.1 | 10.2 | 9.7 | 7.3 |
| May | 12.8 | 13.9 | 16.4 | 10.6 | 13.4 | 11.9 |
| June | 16.5 | 18.5 | 17.8 | 19.5 | 18.1 | 15.0 |
| July | 18.7 | 18.7 | 19.2 | 19.0 | 18.9 | 16.7 |
| average (March - July) | 12.0 | 13.1 | 13.9 | 13.1 | 13.0 | 10.8 |

**Table S-4.** Recorded sum of precipitation during the growth period during field seasons between 2016 and 2019 and calculated means for each month between March and July. The mean sums of precipitation between 1961 and 1990 are listed as baseline. The data was collected one meter above ground by a weather station at the location in Freising. The data was accessed from agro-meteorology web portal of the Bavarian State Institute for Agriculture (Agrarmeteorologie Bayern, 2020).

|  | sum of precipitation [mm] |  |  |  |  |  |
| --- | --- | --- | --- | --- | --- | --- |
|  | 2016 | 2017 | 2018 | 2019 | average<br>(2016 - 2019) | average<br>(1961-1990) |
| **March** | 41.4 | 42.6 | 30.1 | 48.5 | 40.7 | 43.4 |
| **April** | 48.8 | 82.1 | 10.3 | 12.3 | 38.4 | 55.5 |
| **May** | 98.1 | 68.6 | 91.4 | 118.0 | 94.0 | 89.6 |
| **June** | 105.5 | 56.3 | 123.0 | 79.3 | 91.4 | 103.7 |
| **July** | 93.1 | 115.8 | 69.1 | 54.3 | 83.1 | 98.3 |
| **average**<br>**(March - July)** | **77.4** | **73.1** | **65.1** | **62.5** | **69.5** | **78.1** |

**Table S-5:**
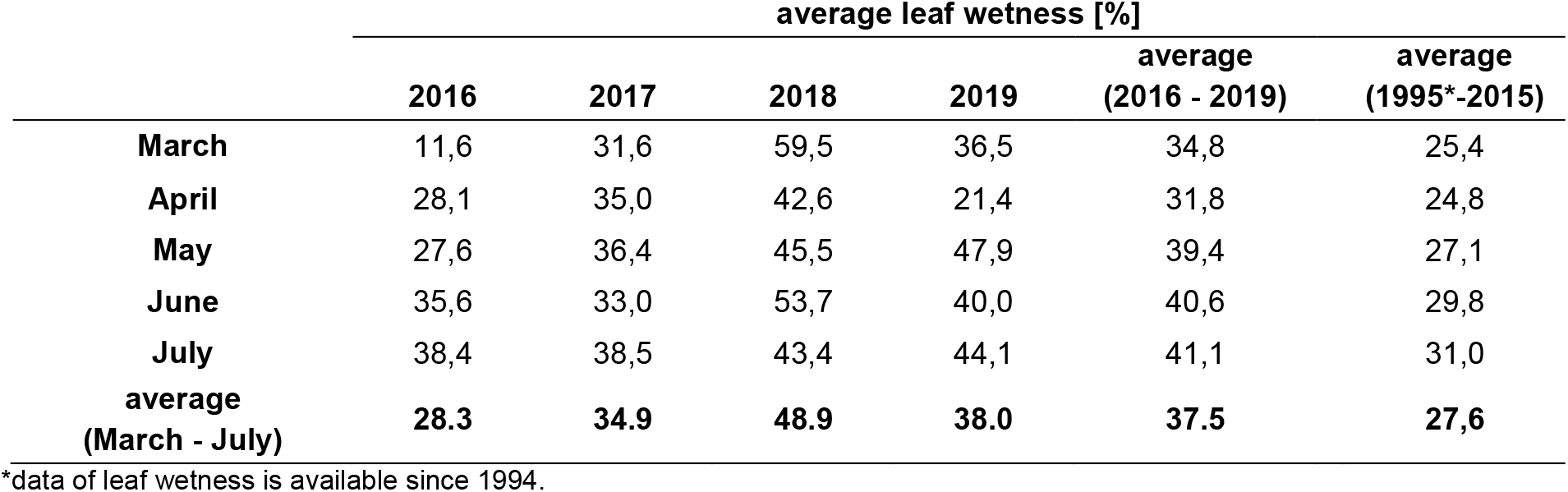
Recorded mean percentage of leaf wetness during the growth period during field seasons between 2016 and 2019 and calculated means for each month between March and July. The means of leaf wetness between 1995* and 2015 are listed as baseline. The data was collected by a weather station at the location in Freising. The data was accessed from agro-meteorology web portal of the Bavarian State Institute for Agriculture (Agrarmeteorologie Bayern, 2020).

**Table S-6.** Recorded average sum of global radiation during the growth period during field seasons between 2016 and 2019 and calculated means for each month between March and July. The mean sums of global radiation between 1995* and 2015 are listed as baseline. The data was collected in two meters above ground by a weather station at the location in Freising. The data was accessed from agro-meteorology web portal of the Bavarian State Institute for Agriculture (Agrarmeteorologie Bayern, 2020).

|  | average sum of global radiation [Wh/m²] |  |  |  |  |  |
| --- | --- | --- | --- | --- | --- | --- |
|  | 2016 | 2017 | 2018 | 2019 | average<br>(2016 - 2019) | average<br>(1995*-2015) |
| **March** | 2841 | 3203 | 2781 | 3170 | 2999 | 2951 |
| **April** | 4103 | 3800 | 5279 | 4842 | 4506 | 4361 |
| **May** | 4926 | 5842 | 5733 | 4492 | 5248 | 5251 |
| **June** | 5363 | 6564 | 5836 | 6851 | 6153 | 5812 |
| **July** | 5561 | 5476 | 6196 | 5745 | 5744 | 5534 |
| **average**<br>**(March - July)** | **4559** | **4977** | **5165** | **5020** | **4930** | **4782** |

\*data of leaf wetness is available since 1994.

